# Mapping vascular network architecture in primate brain using ferumoxytol-weighted laminar MRI

**DOI:** 10.1101/2024.05.16.594068

**Authors:** Joonas A. Autio, Ikko Kimura, Takayuki Ose, Yuki Matsumoto, Masahiro Ohno, Yuta Urushibata, Takuro Ikeda, Matthew F. Glasser, David C. Van Essen, Takuya Hayashi

## Abstract

Mapping the vascular organization of the brain is of great importance across various domains of basic neuroimaging research, diagnostic radiology, and neurology. However, the intricate task of precisely mapping vasculature across brain regions and cortical layers presents formidable challenges, resulting in a limited understanding of neurometabolic factors influencing the brain’s microvasculature. Addressing this gap, our study investigates whole-brain vascular volume using ferumoxytol-weighted laminar-resolution multi-echo gradient-echo imaging in macaque monkeys. We validate the results with published data for vascular densities and compare them with cytoarchitecture, neuron and synaptic densities. The ferumoxytol-induced change in transverse relaxation rate (ΔR_2_*), an indirect proxy measure of cerebral blood volume (CBV), was mapped onto twelve equivolumetric laminar cortical surfaces. Our findings reveal that CBV varies 3-fold across the brain, with the highest vascular volume observed in the inferior colliculus and lowest in the corpus callosum. In the cerebral cortex, CBV is notably high in early primary sensory areas and low in association areas responsible for higher cognitive functions. Classification of CBV into distinct groups unveils extensive replication of translaminar vascular network motifs, suggesting distinct computational energy supply requirements in areas with varying cytoarchitecture types. Regionally, baseline R_2_* and CBV exhibit positive correlations with neuron density and negative correlations with receptor densities. Adjusting image resolution based on the critical sampling frequency of penetrating cortical vessels allows us to delineate approximately 30% of the arterial-venous vessels. Collectively, these results mark significant methodological and conceptual advancements, contributing to the refinement of cerebrovascular MRI. Furthermore, our study establishes a linkage between neurometabolic factors and the vascular network architecture in the primate brain.

**Highlights:** Cortical layer vascular mapping using ferumoxytol-weighted R_2_* MRI

Vascular volume is high in primary sensory areas and low in association areas

Correlation between R_2_* and vascular volume with neuron and receptor densities

Vascularization co-varies with densities of specific interneuron types

## 1 Introduction

The brain’s vascular network plays a crucial role in delivering oxygen, glucose, and other nutrients while clearing metabolic by-products to meet the high energy demands of neural information processing. Understanding the organization of the brain’s vasculature is vital for diagnosing and addressing clinical deficits related to stroke, vascular dementia, and neurological disorders with vascular components (Iadecola, 2013; Sweeney et al., 2018; Toledo et al., 2013). Furthermore, it is essential for advancing the applications of functional MRI (fMRI), as vascular density has implications for statistical power, and the arrangement of large vessels may impose limitations and biases on the spatial accuracy of functional localization. Despite its significance, our knowledge of the vascular network architecture in the primate cerebral cortex remains limited (Duvernoy et al., 1981; Schmid et al., 2019; Weber et al., 2008).

Anatomically, blood flows from the pial vessel network via feeding arteries and arterioles to capillary beds in each cortical layer, ultimately leading via draining veins back to the pial vessel network. The capillary density varies with the rate of oxidative metabolism across cortical layers and exhibits sharp transitions between some cortical areas (Duvernoy et al., 1981; Ji et al., 2021; Zheng et al., 1991). Recent advances in immunolabeling and tissue clearing techniques have enhanced our understanding of brain vascularity in post-mortem mouse brains (Ji et al., 2021; Kirst et al., 2020). These studies have demonstrated heterogeneous vasculature varying in capillary length density 3-fold across brain regions and moderate variation across cortical layers. Still, methodological and analytical challenges have limited quantitative anatomical research in primates to a small number of cortical regions (Harrison et al., 2002; Lauwers et al., 2008; Weber et al., 2008) and quantitative anatomical research requires investigation across a broader range of cortical regions in primates.

Mapping the brain-wide vasculature using MRI faces several challenges due to the intricate nature of the vascular network. One crucial criterion for successful vascular mapping is arterio-venous density, which is necessary to delineate individual large-caliber vessels from microcapillary networks. The combined surface density of intra-cortical feeding arteries and draining veins is about 7 vessels/mm^2^ (Weber et al., 2008). According to the sampling theorem, this implies that the minimal (spatial) sampling frequency is ≈14 voxels/mm^2^ (≈0.26 mm isotropic) imposing stringent image acquisition requirements to critically sample cortical vasculature. Ferumoxytol contrast agent-weighted MRI offers a safe and indirect means to measure relative vascular volume and enhance the visibility of large vessels (Boxerman et al., 1995; Kim et al., 2013; Muehe et al., 2016; Yablonskiy and Haacke, 1994). Compared to clinically used gadolinium-based agents, ferumoxytol’s substantially longer half-life and stronger R_2_* effect allows for higher-resolution and more sensitive vascular volume measurements (Buch et al., 2022), albeit these methodologies are hampered by confounding factors such as vessel orientation relative to the magnetic field (B_0_) direction (Ogawa et al., 1993).

The macaque monkey is an excellent experimental non-human primate model to objectively investigate the MRI resolution and contrast requirements and their limitations for mapping arterio-capillary-venous networks. Quantitative vascular density data is available for a limited number of cortical areas (Weber et al., 2008), providing essential insights for determining vessel-density informed minimum image resolution requirements. Importantly, experiments in macaque monkeys can also help elucidate the neurometabolic factors that shape vascular network architecture. For instance, variations in the cellular composition (Collins et al., 2010), synaptic density (Elston, 2002), receptor distribution (Froudist-Walsh et al., 2023), neural connectivity (Felleman and Van Essen, 1991; N. T. Markov et al., 2014), myelination (Lewis and Essen, 2000), and oxidative metabolism (Sincich et al., 2003) are well-documented, but the relationships between these factors and vascular architecture have only been investigated in a few cortical areas (Borowsky and Collins, 1989; Tsai et al., 2009; Weber et al., 2008).

In this study, we used ferumoxytol contrast agent-weighted 3D multi-echo gradient-echo MRI to investigate vascular heterogeneity in macaque monkey brains. We scaled image resolution to meet specific requirements, aiming to delineate cortical layers and individual vessels. Using this advanced MRI and laminar surface mapping, we then elucidate neuroanatomical factors underlying the heterogenous vasculature. Our analysis reveals insights into translaminar and regional heterogeneities, and signatures of neuroanatomical organization within the macaque cerebral cortex. By addressing these dual objectives—advancing vascular MRI technology and uncovering the neuroanatomical factors shaping cortical vascularity—we contribute to both methodological and conceptual advancements in the field. Our findings offer not only a framework for objectively validating cerebrovascular MRI but also a deeper understanding of how neural and vascular systems intricately interact in the primate cerebral cortices.

## 2 Methods

### 2.1 Data acquisition

Experiments were performed using a 3T MRI scanner (MAGNETOM Prisma, Siemens, Erlangen, Germany) equipped with 80 mT/m gradients (XR 80/200 gradient system with slew rate 200 T/m/s), a 2-channel B_1_ transmit array (TimTX TrueForm) and a custom-made 24-channel coil for the macaque brain (Autio et al., 2020). The animal experiments were conducted in accordance with the institutional guidelines for animal experiments, and animals were maintained and handled in accordance with the policies for the conduct of animal experiments in research institution (MEXT, Japan, Tokyo) and the Guide for the Care and Use of Laboratory Animals of the Institute of Laboratory Animal Resources (ILAR; Washington, DC, USA). All animal procedures were approved by the Animal Care and Use Committee of the Kobe Institute of RIKEN (MA2008-03-11).

#### 2.1.1 Anesthesia protocol

Macaque monkeys (*Macaca mulatta*, weight range 7.4 – 8.4 kg, age range 4-6 years, N = 4) were initially sedated with intramuscular injection of atropine sulfate (20 μg/kg), dexmedetomidine (4.5 µg/kg) and ketamine (6 mg/kg). A catheter was inserted into the caudal artery for blood-gas sampling, and endotracheal intubation was performed for steady controlled ventilation using an anesthetic ventilator (Cato, Drager, Germany). End-tidal carbon dioxide was monitored and used to adjust ventilation rate (0.2 to 0.3 Hz) and end-tidal volume. After the animal was fixed in an animal holder, anesthesia was maintained using 1.0% isoflurane via a calibrated vaporizer with a mixture of air 0.75 L/min and O2 0.1 L/min. Animals were warmed with a blanket and water circulation bed and their rectal temperature (1030, SA Instruments Inc., NY, USA), peripheral oxygen saturation and heart rate (7500FO, NONIN Medical Inc., MN, USA) were monitored throughout experiments.

#### 2.1.2 Structural acquisition protocol

T1w images were acquired using a 3D Magnetization Prepared Rapid Acquisition Gradient Echo (MPRAGE) sequence (0.5 mm isotropic, FOV 128×128×112 mm, matrix 256×256, slices per slab 224, coronal orientation, readout direction of inferior (I) to superior (S), phase oversampling 15%, averages 3, TR 2200 ms, TE 2.2 ms, TI 900 ms, flip-angle 8.3°, bandwidth 270 Hz/pixel, no fat suppression, GRAPPA 2, turbo factor 224 and pre-scan normalization). T2w images were acquired using a Sampling Perfection with Application optimized Contrast using different angle Evolutions (SPACE) sequence (0.5 mm isotropic, FOV 128×128×112 mm, matrix 256×256, slice per slab 224, coronal orientation, readout direction I to S, phase oversampling 15%, TR 3200 ms, TE 562 ms, bandwidth 723 Hz/pixel, no fat suppression, GRAPPA 2, turbo factor 314, echo train length 1201 ms and pre-scan normalization) (Autio et al., 2021, 2020)

In a separate imaging session, additional high-resolution structural images were acquired (Autio et al., 2024). T1w images were acquired using a 3D Magnetization Prepared Rapid Acquisition Gradient Echo (MPRAGE) sequence (0.32mm isotropic, FOV 123×123×123 mm, matrix 384×384, slices per slab 256, sagittal orientation, readout direction FH, averages 12-15, TR 2200 ms, TE 3 ms, TI 900 ms, flip-angle 8°, bandwidth 200 Hz/pixel, no fat suppression, GRAPPA 2, reference lines PE 32, turbo factor 224, averages 12-15, and pre-scan normalization). T2w images were acquired using a Sampling Perfection with Application optimized Contrast using different angle Evolutions and Fluid-Attenuated Inversion Recovery (SPACE-FLAIR) sequence (0.32mm isotropic, FOV 123×123×123 mm, matrix 384×384, slice per slab 256, sagittal orientation, readout direction FH, TR 5000ms, TE 397 ms, TI 1800 ms, bandwidth 420 Hz/pixel, no fat suppression, GRAPPA 2, reference lines PE 32, turbo factor 188, echo train duration 933 ms, averages 6-7 and pre-scan normalization). The total acquisition time for structural scans was ≈3h.

#### 2.1.3 Quantitative transverse relaxation rate acquisition protocol

Data was acquired before and after (12 mg/kg) the intravascular ferumoxytol (Feraheme, ferumoxytol AMAG Pharmaceuticals Inc, Waltham, MA, USA) injection using gradient- and RF-spoiled 3D multi-echo gradient-echo acquisition (0.32 mm isotropic, FOV 103×103×82 mm, matrix 320×320, slices per slab 256, sagittal orientation, bipolar read-out mode, elliptical scanning, no partial Fourier, ten equidistant TEs, first TE (TE1)=3.4 ms, time between echoes (ΔTE) 2.4 ms, TR 33 ms, FA 13° (corresponding to Ernst angle of gray matter; median T_1_=1370 ms), bandwidth 500 Hz/pixel (fat-water shift one voxel), scan duration 20 min, GRAPPA 2, reference lines 32, and pre-scan normalization). The total acquisition time before and after ferumoxytol injection were 40 and 100 min, respectively.

#### 2.1.4 Vessel-density informed data acquisition protocol

To investigate the periodicity of the penetrating large vessel network, we performed auxiliary ferumoxytol-weighted experiments using image resolution adjusted to satisfy critical (spatial) sampling frequency (14 voxels/mm^2^ ≈0.26 mm isotropic) of intra-cortical vessels (7 vessels/mm^2^; Weber et al., 2008). The original gradient- and RF-spoiled 3D multi-echo gradient-echo product sequence, however, did not allow sufficient matrix size to satisfy the critical sampling frequency of penetrating vessels. To achieve the target resolution, the sequence was customized by easing the matrix size limitations (by Y.U.). Using the customized sequence, we performed experiments at 0.25 (N=1) and 0.23 mm (N=2) isotropic spatial resolution. Scan #1: (FOV 104×104×80 mm, matrix 416×416, slices per slab 320, sagittal orientation, bipolar read-out mode, elliptical scanning, no partial Fourier, three TEs 6, 10 and 14 ms, TR 22 ms, FA 11°, bandwidth 260 Hz/pixel (fat-water shift 1.6 voxels), scan duration 21 min, GRAPPA 2, reference lines 32 and pre-scan normalization). Scans #2-3: (FOV 103×103×81 mm, matrix 448×448, slices per slab 352, sagittal orientation, bipolar read-out mode, elliptical scanning, no partial Fourier, three TEs 5, 9 and 13 ms, TR 23 ms, FA 11°, bandwidth 340 Hz/pixel (fat-water shift 1.2 voxels), scan duration 25 min, GRAPPA 2, reference lines 32 and pre-scan normalization). The total acquisition time was 150 min.

### 2.2 Data analysis

Data analysis utilized a version of the HCP pipelines customized specifically for use with non-human primates (https://github.com/Washington-University/NHPPipelines) (Autio et al., 2020; Glasser et al., 2013; Hayashi et al., 2021).

#### 2.2.1 Structural image processing

PreFreeSurfer pipeline registered structural T1w and T2w images into an anterior-posterior commissural (AC-PC) alignment using a rigid body transformation, non-brain structures were removed, T2w and T1w images were aligned using boundary based registration (Greve and Fischl, 2009), and corrected for signal intensity inhomogeneity using B_1_-bias field estimate (Glasser et al., 2013). Next, data was transformed into a standard macaque atlas by 12-parameter affine and nonlinear volume registration using FLIRT and FNIRT FSL tools (Jenkinson et al., 2002).

FreeSurferNHP pipeline was used to reconstruct the cortical surfaces using FreeSurfer v6.0.0-HCP. This process included conversion of data in native AC-PC space to a ‘fake’ space with 1-mm isotropic resolution in volume with a matrix of 256 in all directions, intensity correction, segmentation of the brain into cortex and subcortical structures, reconstruction of the white and pial surfaces and estimation of cortical folding maps and thickness. The intensity correction was performed using FMRIB’s Automated Segmentation Tool (FAST) (Zhang et al., 2001). The white matter segmentation was fine-tuned by filling a white matter skeleton to accurately estimate white surface around the blade-like thin white matter particularly in the anterior temporal and occipital lobe (Autio et al., 2020). After the white surface was estimated, the pial surface was initially estimated by using intensity normalized T1w image and then estimated using the T2w image to help exclude dura (Glasser et al., 2013).

The PostFreeSurfer pipeline transformed anatomical volumes and cortical surfaces into the Yerkes19 standard space, performed surface registration using folding information via MSMSulc (Robinson et al., 2018, 2014), generated mid-thickness, inflated and very inflated surfaces, as well as the myelin map from the T1w/T2w ratio on the mid-thickness surface. The volume to surface mapping of the T1w/T2w ratio was carried out using a ‘myelin-style’ mapping (Glasser and Van Essen, 2011), in which a cortical ribbon mask and a metric of cortical thickness were used, weighting voxels closer to the midthickness surface. Voxel weighting was done with a Gaussian kernel of 2 mm FWHM, corresponding to the mean cortical thickness of macaque. For quality control, the myelin maps were visualized and potential FreeSurfer errors in pial or WM surface placement were identified. The errors were manually corrected by editing wm.mgz and by repositioning and smoothing the surfaces using FreeSurfer 7.1, the curvature, thickness, and surface area on each vertex were recalculated, and then PostFreeSurfer pipeline was applied again and T1w/T2w was visually inspected for quality control.

Twelve cortical laminar surfaces were generated based on equivolume model (Autio et al., 2024; Van Essen and Maunsell, 1980) using the Workbench command ‘-surface-cortex-layer’ and the native pial and white surface meshes in subject’s AC-PC space. Throughout the text, the equivolumetric layers (ELs) are referred to as EL1a (adjacent to the pial surface), EL1b, EL2a, EL2b,… and EL6b (adjacent to the white matter surface). This nomenclature is intended to ease but also distinguish comparison between anatomically determined cortical layers which vary in thickness. Anatomical layers are referred to using roman numerals (e.g. Ia, Ib, Ic, IIa,… and VIb). To assess the vascularity on the white matter and pial surfaces, additional layers were generated underneath and just above the gray matter in the superficial white matter and pial surface, respectively. Surface models and data were resampled to a high-resolution 164k mesh (per hemisphere).

#### 2.2.2 Quantitative multi-echo gradient-echo data processing

The original 3D multi-echo gradient-echo images were upsampled to 0.25 or 0.15 mm spatial resolution for the data with spatial resolution 0.32 or 0.23 and 0.26, respectively; and transformed using cubic-spline to the subject’s AC-PC space using a rigid body transformation. Pre- and post-ferumoxytol runs (two and six, respectively) were averaged and R_2_*-fitting procedure was performed on multi-TE images with ordinary least squares method in the in vivo histology using MRI (hMRI) Toolbox (Tabelow et al., 2019). The baseline (pre-ferumoxytol) R_2_* was subtracted from the post-ferumoxytol R_2_* maps to calculate ferumoxytol induced change in ΔR_2_* (Boxerman et al., 1995). Subcortical region-of-interests (ROIs; thalamus, striatum, cerebellum, hippocampus, inferior colliculus and corpus callosum) were manually drawn whilst avoiding large vessels using T1w image as a reference. The quantitative R_2_* and ΔR_2_* maps were mapped in the twelve native laminar mesh surfaces using the Workbench command ‘-volume-to-surface-mapping’ using a ribbon-constrained algorithm. MSMSulc surface registration was applied, the data was resampled to Mac25Rhesus reference sulcus template using ADAP_BARY_AREA with vertex area correction, and left and right hemispheres were combined into a CIFTI file.

Since large-caliber pial vessels run along cortical surface, large penetrating vessels are mainly oriented normal to the cortical surface and the capillary network may be orientated more random to the cortical surface (Ji et al., 2021; Reina-De La Torre et al., 1998), the orientation of the cerebral cortex relative to the direction of static magnetic field (B_0_) may bias the assessment of R_2_* and ΔR_2_* (Bolan et al., 2006; Lee et al., 2011; Ogawa et al., 1993; Viessmann et al., 2019; Yablonskiy and Haacke, 1994). Because the brain R_2_* measures are primarily determined by extravascular MR signal, we may assume that

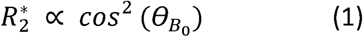

where Θ is the angle between normal of the cortex relative to the direction of B_0_. Each vertices Θ was determined in the subject’s original MRI space. The (Pearson’s) correlation coefficient between Eq. 1 and with R_2_* and ΔR_2_* was estimated in each EL. To remove orientation bias, in Eq. 1 was regressed out from each laminar R_2_* and ΔR_2_* surface map.

To examine repetitive patterns in the vascularity, the R_2_* and ΔR_2_* laminar profiles were parcellated using the M132 Lyon Macaque brain atlas (N. T. Markov et al., 2014). Because some of the M132 atlas cortical parcels exhibited a degree of laminar inhomogeneity due to artifacts (e.g., areas adjacent to major sinuses, large vessels penetrating to white matter and FreeSurfer errors in surface placement), median values were assigned to each parcel in each EL. The effect of blood accumulation in large feeding arteries and draining veins toward the superficial layers was estimated using linear model and regressed out from the parcellated ΔR_2_* maps. Subjects (including a single test-retest data set), ELs and hemispheres were combined (5×12×2=120) and hierarchical clustering was applied to parcels using Ward’s method. A dendrogram was used to determine the number of clusters.

To explore sharp transitions in cortical vascularization, each B_0_ orientation corrected ΔR_2_* EL was smoothed using a factor of 1.2 mm. The smoothing factor was twice the average distance of draining veins (Weber et al., 2008). Then, gradient-ridges were calculated using maps using wb_command -cifti-gradient for each EL. The resulting gradients were then cross-referenced with potential FreeSurfer surface error displacements and when required manual corrections (wm.mgz and using reposition surface in the FreeView 3.0) were performed to the FreeSurfer segmentations.

#### 2.2.3 Vessel detection

To improve contrast-to-noise ratio for vessel detection, multi-echo gradient-echo images were aligned in the native AC-PC space and averaged across runs. Vessels were identified using the Frangi “vesselness” filter which enhances the vessel/ridge-like structures in 3D image using hessian eigenvalues (Avadiappan et al., 2020; Frangi et al., 1998). Volume images in the native AC-PC space were also non-linearly transformed into a standard “SpecMac25Rhesus” atlas (Hayashi et al., 2021).

To facilitate visualization of the pial vessel network, low-frequency fluctuations were removed by subtracting extensively smoothed versions of the post-ferumoxytol TE-averaged EL surface maps. To detect vessels running parallel to the cortical surface, continuous signal drop-outs were clustered along ELs using wb_command -cifti-find-clusters with a criterion of a 0.5 mm^2^ minimum cluster area and visually determined intensity threshold.

To enable surface detection of penetrating vessels, an ultra high-resolution 656k cortical surface mesh was generated using wb_command -surface-create-sphere resulting in an average 0.022 vertex surface area, approximately half the isotropic 0.23 mm voxel (face) surface area (0.053 mm^2^). Then, TE-averaged multi-echo gradient-echo images and Frangi-filtered vessel images were mapped to twelve native mesh laminar surfaces in the subject’s physical space. To detect penetrating vessels oriented perpendicular to the laminar surfaces, localized signal drop-outs were visualized using wb_command -cifti-gradient and their central locations were identified by detecting the local minima using wb_command -find-extrema.

The number of vessels in V1 was estimated using the M132 parcellation as a reference (Markov et al., 2014). In the native space, the surface area of each vertex was determined using wb_command -surface-vertex-areas. Then, the surface-area map was transferred into atlas space and the area of V1 was determined using M132 areal atlas. The V1 vessel density was determined by dividing the number of vessels (by a conservative estimate using Frangi-filtering and a liberal estimate using local minima in gradient) by surface-area. These MRI estimates were then compared with histological large vessel densities in the V1 (Zheng et al., 1991, Weber et al., 2008).

To determine the periodicity of the cortical arterio-venous networks, non-uniformly sampled Lomb-Scargle geodesic periodogram analysis (Matlab Signal Processing Toolbox, The MathWorks Inc., US) was performed on the spatially low-frequency filtered twelve native mesh ELs in the subject’s physical space. The analysis was limited to the closest 2000 geodesic vertices within 20 mm geodesic distance of manually selected vertices in V1. Geodesic distance between vertices was calculated with wb_command -surface-geodesic-distance in each EL. Periodograms were binarized with an equidistant interval (=0.05 1/mm) up to 10 1/mm, and then 95% confidence interval of the mean of magnitude was estimated from bootstrap and compared across cortical laminae.

#### 2.2.4 Comparison with histological datasets

In V1, translaminar ΔR_2_* was compared with cytochrome oxidase (CO) activity, capillary and large vessel volume fractions (Weber et al., 2008) These ground-truth measures were estimated from Weber and colleagues Figure 4. Each measure was peak normalized, so that values ranged between zero and one, to compare different contrasts across the cortical layers.

To investigate the correspondence between regional variation in cerebrovascular volume and heterogeneous neuron density, we used the 42 Vanderbilt tissue sections covering the entire macaque cerebral cortex (Collins et al. 2010) available from the Brain Analysis Library of Spatial Maps and Atlases (BALSA) database (Froudist-Walsh et al., 2023, 2021; Van Essen et al., 2017). These sections were processed using the isotropic fractionator method to estimate neuron densities (Collins et al., 2010). The sections were used to parcellate R_2_* and ΔR_2_* and these were then compared with neuron density using Pearson’s correlation coefficient. To compare neuron and total receptor densities with R_2_* and ΔR_2_*, we also applied the Julich Macaque Brain Atlas for parcellation (Froudist-Walsh et al., 2023). The parcellated neuron and total receptor densities were used in linear regression model to predict R_2_* and ΔR_2_* across ELs and the resulting T-values were then threshold at significance level (*p*<0.05, Bonferroni-corrected).

Relation between cerebrovascular volume and parvalbumin and calretinin positive interneurons, collated from multiple studies and ascribed to M132 macaque atlas by Burt and colleagues (Burt et al., 2018; Condé et al., 1994; Gabbott and Bacon, 1996; Kondo et al., 1999), were compared across ELs using Pearson’s correlation coefficient. The resulting p-values were Bonferroni-corrected for the number of layers and contrasts.

The number of dendritic spines (putative excitatory inputs) and dendrite tree length size were also obtained from BALSA (Elston, 2007; Froudist-Walsh et al., 2023, 2021). The region-of-interests, described in Elston 2007 and plotted on the Yerkes surface by Froudist-Walsh and colleagues, were used to obtain median value of R_2_* and ΔR_2_* and these were then compared with the number of dendritic spines and dendrite tree length size using Pearson’s correlation coefficient.

## 3 Results

### 3.1 Laminar R_2_* and ferumoxytol induced ΔR_2_* MRI in macaque cerebral cortex

Figure 1 displays representative gradient-echo images before (Fig. 1A) and after (Fig. 1B) intravascular ferumoxytol injection (N=1). The ferumoxytol effectively reversed the signal-intensity contrast between gray matter and white matter while enhancing the visibility of large vessels, as expected. For quantitative assessment, R_2_* values were estimated from multi-echo gradient-echo images acquired both before and after the administration of ferumoxytol contrast agent (Table 1). Subsequently, the baseline R_2_* and ΔR_2_*, an indirect proxy measure of CBV (Boxerman et al., 1995), volume maps for each subject were mapped onto the twelve native equivolumetric layers (ELs) (Fig. 1C). Each vertex was then corrected for normal of the cortex relative to B_0_ direction (Supp. Fig. 1A-C). Surface maps for each subject were registered onto a Mac25Rhesus average surface using cortical curvature landmarks and then averaged across the subjects (Fig. 1D, E). Around cortical midthickness, the distribution of R_2_*, an aggregate measure for ferritin-bound iron, myelin content and venous oxygenation levels (Langkammer et al., 2012), resembled the spatial pattern of ΔR_2_* vascular volume. However, across cortical layers, these measures exhibited reversed patterns: R_2_* increased toward the white matter surface whereas ΔR_2_* decreased (Fig. 1D, G).

**Figure 1.**
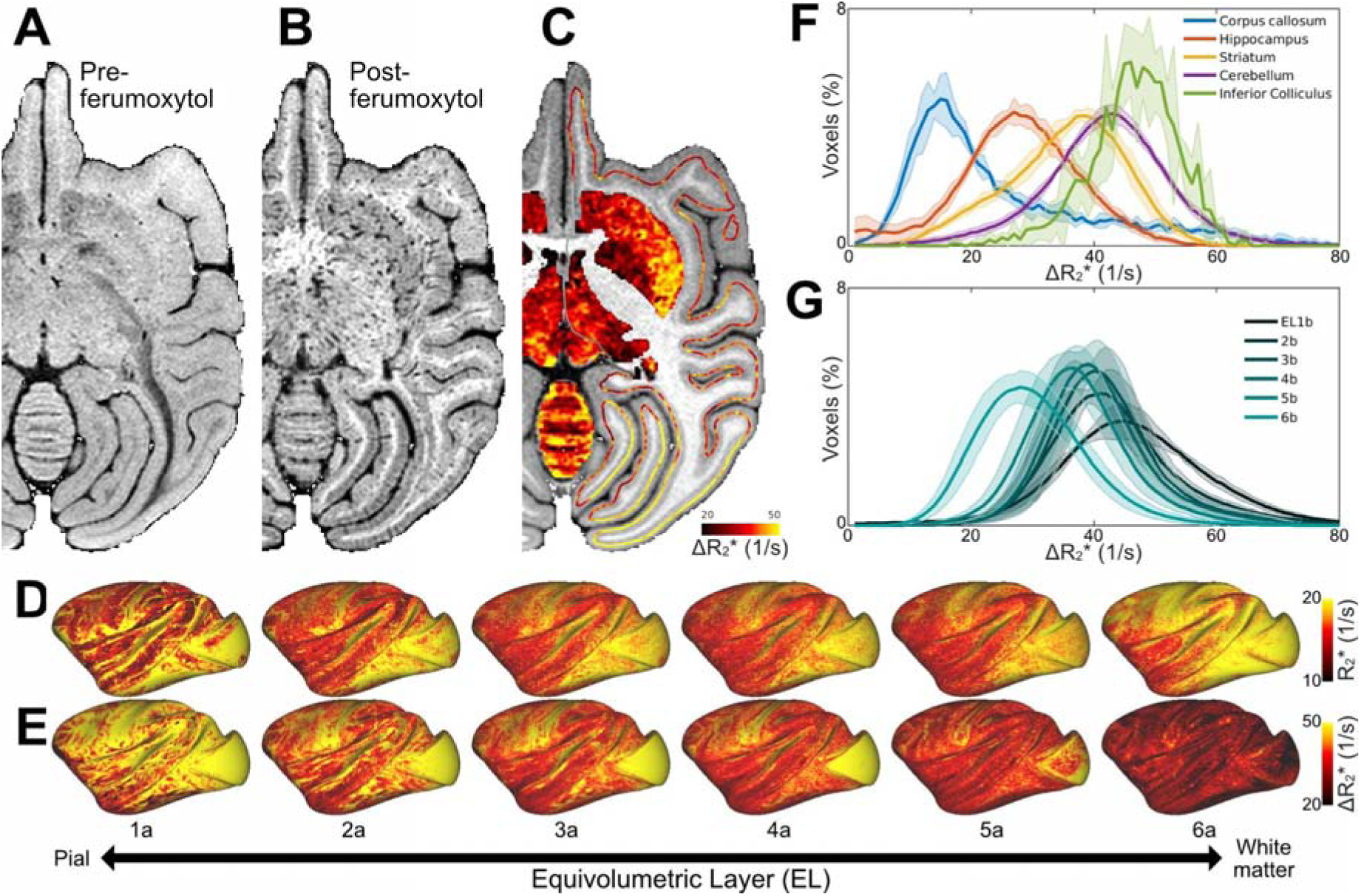
Ferumoxytol-weighted MRI reveals heterogeneous vascularity in the macaque brain. Representative 3D gradient-echo images **(A)** before and **(B)** after the ferumoxytol contrast agent injection. **(C)** Ferumoxytol induced change in transverse relaxation rate (ΔR_2_*) displayed on subcortical gray matter and cortical midthickness surface contour (N=1). Average **(D)** pre-ferumoxytol R_2_*and **(E)** ΔR_2_* equivolumetric layers (ELs; N=4). **(F)** Histograms in selected brain regions and **(G)** ELs. Solid lines and shadow indicate mean and standard deviation (N=4), respectively. Abbreviations: CAU: Caudate nucleus; PUT: Putamen; TH: Thalamus; IC: Inferior colliculus; CBX: Cerebellar cortex.

**Table 1.** Estimated transverse relaxation rate (R_2_*) before and after injection of ferumoxytol contrast agent. Values are mean (std) (N=4). Abbreviations: WM white matter.

| | $R_2^*$ [$s^{-1}$] | Ferumoxytol $R_2^*$ [$s^{-1}$] | $\Delta R_2^*$ [$s^{-1}$] | Vascular volume (%) |
| --- | --- | --- | --- | --- |
| **Cortex** | $17.5 \pm 0.6$ | $56.4 \pm 1.3$ | $38.9 \pm 1.4$ | 2.0 - 2.2<sup>a</sup> |
| **WM** | $21.6 \pm 0.6$ | $45.1 \pm 1.0$ | $23.5 \pm 0.7$ | 0.9 - 1.2<sup>a</sup> |
| **Ratio** | $0.8 \pm 0.1$ | $1.3 \pm 0.1$ | $1.7 \pm 0.1$ | 1.8 - 2.1<sup>a</sup> |

References: <sup>a</sup>Weber et al., (2008).

To explore heterogeneous brain vascularity, we investigated ΔR_2_* in selected subcortical regions (Fig. 1F). We found the highest CBV in the inferior colliculus, an early auditory nucleus, and the lowest in the corpus callosum. Overall, the relative blood volume variations among the investigated subcortical regions were comparable to those reported in mice (Kirst et al., 2020).

Adjacent to the pial surface, large vessels exhibited notable signal-loss in the superficial gray matter. To visualize the pial vessel network, we removed very low-frequency components from TE-averaged post-ferumoxytol signal-intensity maps and identified continuous signal drop-outs along ELs using clustering. Within the most superficial layers (e.g. EL1a-2a), this analysis revealed an extensive arterio-venous pial vessel network spanning almost the entire cortical surface (Fig. 2A). We attempted to delineate pial arteries and veins using pre-contrast R_2_*-values; however, due to the ‘blooming’ effect of ferumoxytol (Buch et al., 2022) distinguishing adjacent large-caliber vessels was difficult to differentiate with high confidence. Additionally, the continuity of the pial vessel network may also have been influenced by veins crossing the sulci (Duverney et al., 1981). Pial vessel network was consistently observed across subjects (Supp. Fig. 2A), although the precise locations of vessels did vary across subjects. In contrast, in the middle and deep cortical layers (EL2b-6b) continuous signal-dropout clusters were largely absent demonstrating that the influence of the large pial vessels was minimal in these layers (Supp. Fig. 2B).

**Figure 2.**
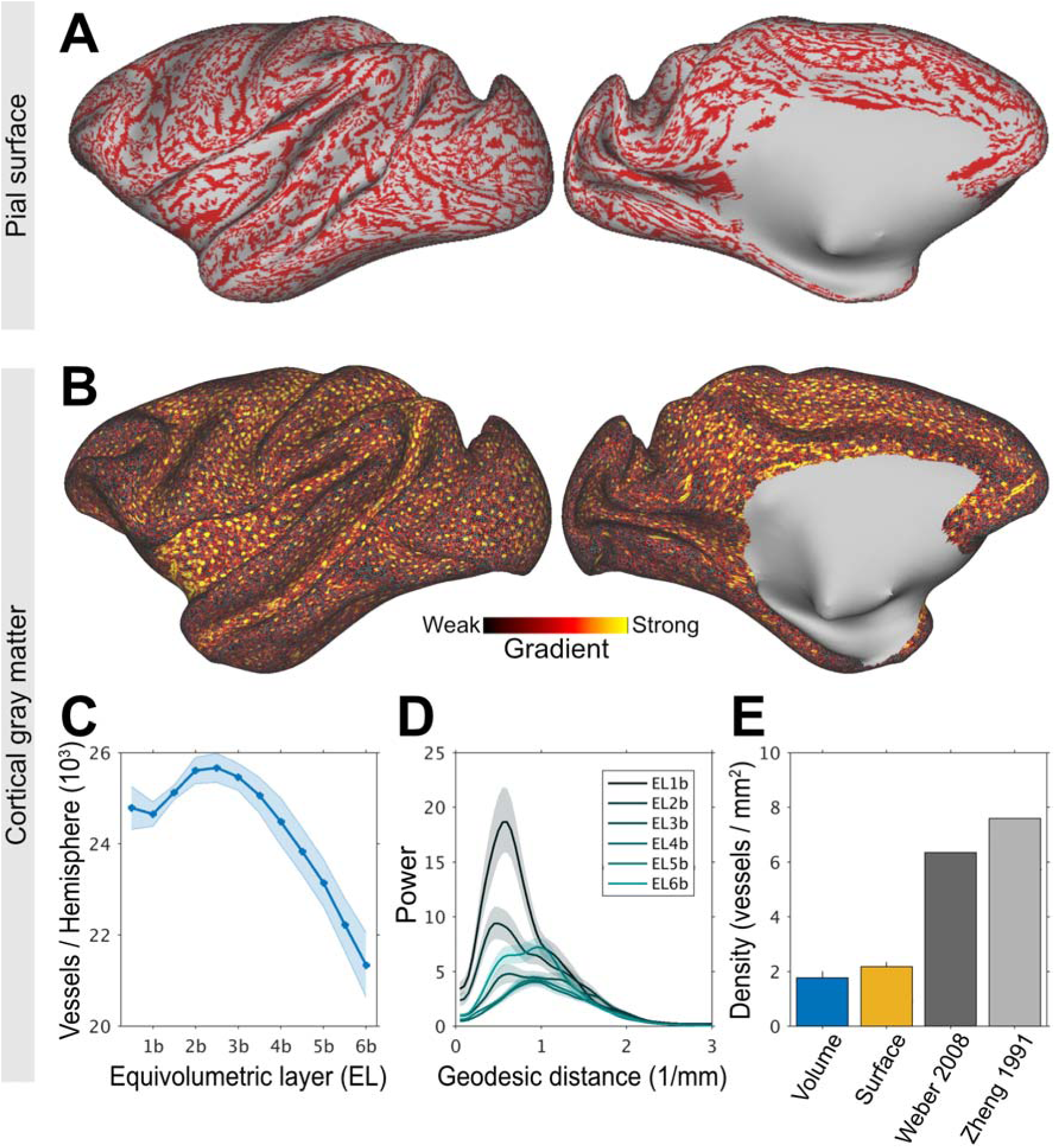
Charting large-caliber vessel networks in the cerebral cortex. **(A)** Ferumoxytol-weighted MRI reveals a continuous pial vessel network running along the cortical surface. Note that the large vessels branch into smaller pial vessels. **(B)** Cortical surface mapping of intra-cortical vessels. Vessels were identified using high-frequency gradients (red-yellow colors) and each blue dot indicates the vessel’s central location. Representative equivolumetric layer (EL) 4a is displayed on a 656k surface mesh. **(C)** Number of penetrating vessels across ELs per hemisphere. Solid lines and shadow show mean and standard deviation across TEs (N=1). **(D)** Non-uniformly sampled Lomb-Scargle geodesic-distance periodogram. The vessels exhibit a peak frequency at about 0.6 1/mm reflecting the frequency of large-caliber vessels. **(E)** Comparison of vessel density in V1 determined using MRI (current study) and ‘ground-truth’ anatomy. In volume space, the density of vessels was estimated using Frangi-filter whereas in the surface mesh the density was estimated using local minima. These are compared to the density of penetrating vessels with a diameter of 20-50 μm (Zheng et al., 1991) and the density of feeding arterial and draining veins evaluated using fluorescence microscopy (Weber et al., 2008).

To visualize the intra-cortical vessel network, we next performed ferumoxytol-weighted experiments with isotropic image resolution of 0.23 mm adjusted below to the critical (spatial) sampling frequency of large penetrating vessels (19 voxels/mm^2^ vs 7 vessels/mm^2^) (Zheng et al., 1991; Weber et al., 2008). The post-ferumoxytol signal-intensity maps (Supp. Fig. 3A) were used to identify vessels in volume space using the Frangi filter (Supp. Fig. 3B) and in the cortical surface by calculating sharp gradients and determining their local minima (Supp. Fig. 3C). Local minima, however, by mathematical definition can capture 1 vessel per 7 vertices (each vertex contains six neighbors). To address this limitation, we generated an ultra-high cortical surface mesh (656k) with an average vertex area of 0.022 ± 0.012 mm^2^ (1st-99th percentile range: 0.006-0.065 mm^2^) (Fig. 2B). Within the cortical gray matter, we identified an average of 24,000 ± 2,000 penetrating vessels per hemisphere (Fig. 2B, C; for all ELs see Supp. Fig. 4). In V1, we found 1.9-2.2 vessels/mm^2^ using Frangi filter and surface vessel detection, respectively (Fig. 2E). This vessel density corresponds to about 30% of the anatomical ground-truth (Zheng et al., 1991, Weber et al., 2008).

To corroborate the periodicity of the cerebrovascular network, we next applied a non-uniformly sampled lomb-scargle geodesic periodogram analysis on the signal-intensity averaged native ELs (Fig. 2D). The periodograms revealed dominant periodicity at spatial frequency of ≈0.6 1/mm in the most superficial ELs, likely reflecting the presence of large-caliber vessels in the pial network. In the middle ELs, bimodal distribution was observed with peaks at around 0.6 and 1.2 1/mm. In the deep ELs, peak power occurred at a shorter distance (≈1 1/mm), potentially indicative of the large arteries supplying the white matter. These findings underscore the substantial variation in vascular organization across the cortical layers.

To explore areal differences in translaminar features, we next parcellated the dense R_2_* and ΔR_2_* maps using M132 cortical atlas (Fig. 2A-D). To mitigate bias resulting from undersampling the large-caliber vessels, median parcel values were used for parcellation, ΔR_2_* profiles were detrended across ELs and then averaged across subjects. In the EL4b, which approximately corresponds to the location of histologically defined thalamic input L4c, the primary visual area exhibited larger vascular volume in comparison to surrounding cortical areas (Fig. 3B, D). Within the visual system, inspection of laminar profiles revealed distinct features.

**Figure 3.**
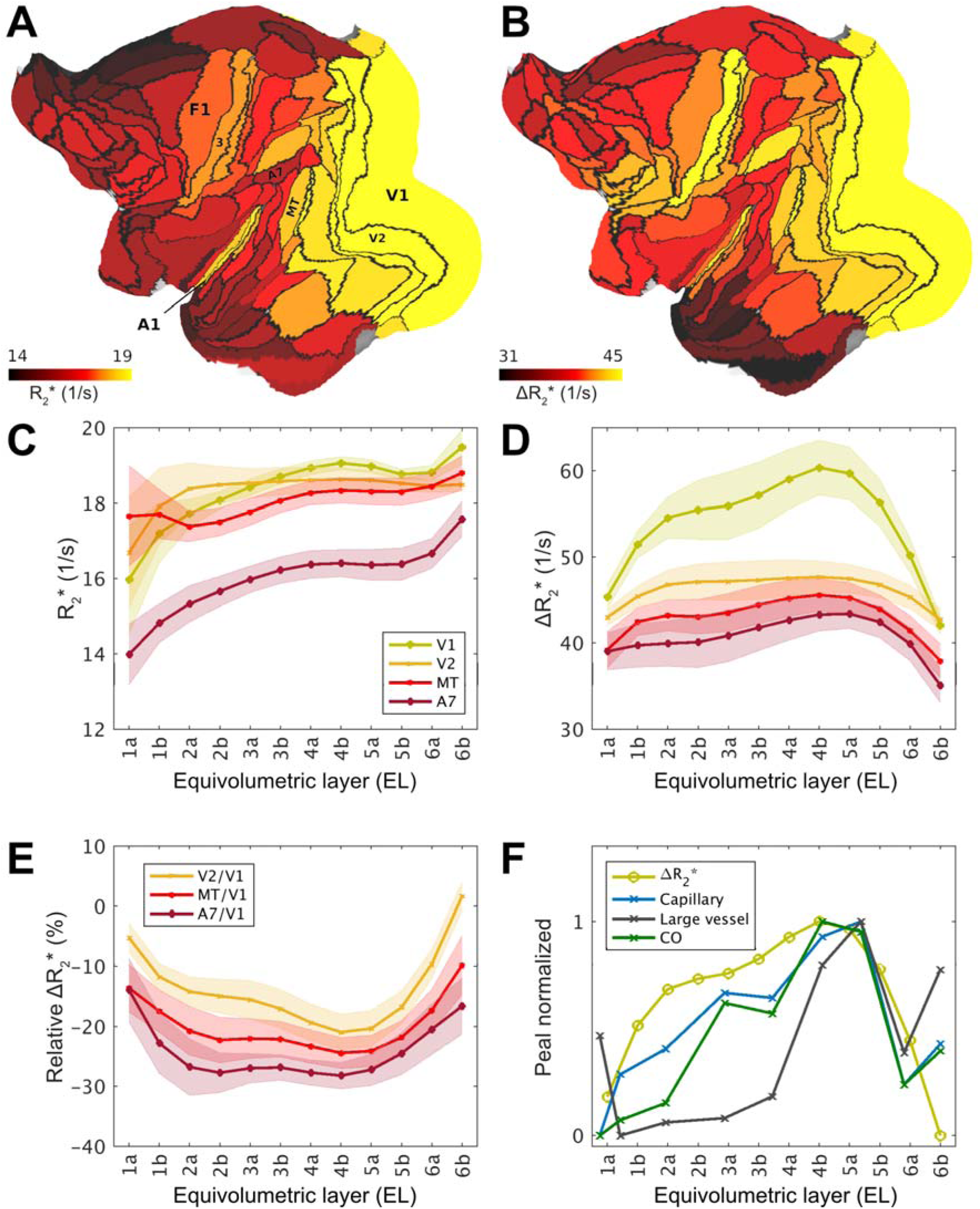
Exemplar laminar profiles of transverse relaxation rate (R_2_*) and ferumoxytol induced change in R_2_ (ΔR_2_*) in the macaque cerebral cortex. **(A)** Exemplar equivolumetric layer 4b (EL4b) R_2_* and **(B)** ΔR_2_* displayed on cortical flat-map. Note that primary sensory areas (e.g. V1, A1, area 3) and association areas exhibit high and low ΔR_2_*, respectively. **(C, D)** Exemplar laminar profiles from visual cortical areas. Solid lines and shadow show mean and standard deviation across hemispheres, respectively. **(E)** Laminar ΔR_2_* profiles relative to the V1. Solid lines and shadow show mean and inter-subject standard deviation. **(F)** Peak-normalized ΔR_2_* profile compared with anatomical ground-truth in V1 (Weber et al., 2008). Cytochrome-c oxidase (CO) activity, capillary and large vessel volume fractions were estimated from their Figure 4. Abbreviations: A1: primary auditory cortex; A7 Brodmann area 7; MT middle temporal area; V1: primary visual cortex; V2: secondary visual cortex; 3: primary somatosensory cortex; 4: primary motor cortex.

Since the ΔR_2_* is an indirect proxy measure of vascular volume (Boxerman et al., 1995), we next sought to validate the noninvasive laminar ΔR_2_* maps with respect to quantitative histological assessment of vascular properties in the macaque visual cortex (Weber et al., 2008). In V1 we found that ΔR_2_* more closely resembled microcapillary and oxidative metabolism rather than large vessel volume fraction (Fig. 3F), albeit we could not identify the vascularity peak in L6 potentially due resolution limitations. Moreover, the V2/V1 ΔR_2_* ratio in EL4b (79% ± 5%) (Fig. 3E) was also in excellent agreement with the previous reports of capillary volume (Zheng et al., 1991; Weber et al., 2008). Finally, the average ΔR_2_* ratio between V1 gray matter and the underlying white matter (2.2 ± 0.1) was also close to the histological assessment (1.8-2.1). Taken together, we found comparable relative variations in vascular volume with anatomical ‘ground-truth’ substantiating the validity of our noninvasive methodology.

### 3.2 Variations in cerebrovascular network architecture reveal inter-areal boundaries

Since cellular composition (Collins et al., 2010) and oxidative metabolism (Sincich et al., 2003) are known to exhibit sharp transitions between cortical areas, we next tested the hypothesis whether the variations in vascular network architecture may also reveal inter-areal boundaries (Zheng et al., 1991). To address this question, we calculated the gradient ridges of ΔR_2_* in each EL (Fig. 1E). Due to the strong cortical contrast (ΔR_2_* = 39 ± 2 ms), the resulting gradients were notably strong and revealed several sharp transitions (Fig. 5A, B). A particularly strong gradient was observed at the boundary between V1/V2 at EL4b (Fig. 3D and Fig. 4A, B), attributable to the relatively large capillary density difference between the areas (Fig. 3E) (Duverney 1981; Zheng et al., 1991; Weber et al., 2008). We also found a sharp ΔR_2_* transition between the primary sensory cortex (area 3) and the primary motor cortex (area 4), in line with histological evaluation of capillary density in humans (Duverney 1981). Moreover, we discovered a sharp ΔR_2_* transition between area 3 and the secondary sensory area (Brodmann area 2). The estimated area boundary locations were supported by comparison of cortical area boundaries as defined in the M132 atlas (Fig. 4A,B) (N. T. Markov et al., 2014). The auditory cortex exhibited also relatively high ΔR_2_*, however, the gradient-ridges were less distinguishable in this region. Multiple layer-specific gradient-ridges were also observed, albeit these were weaker in magnitude and more challenging to delineate.

**Figure 4.**
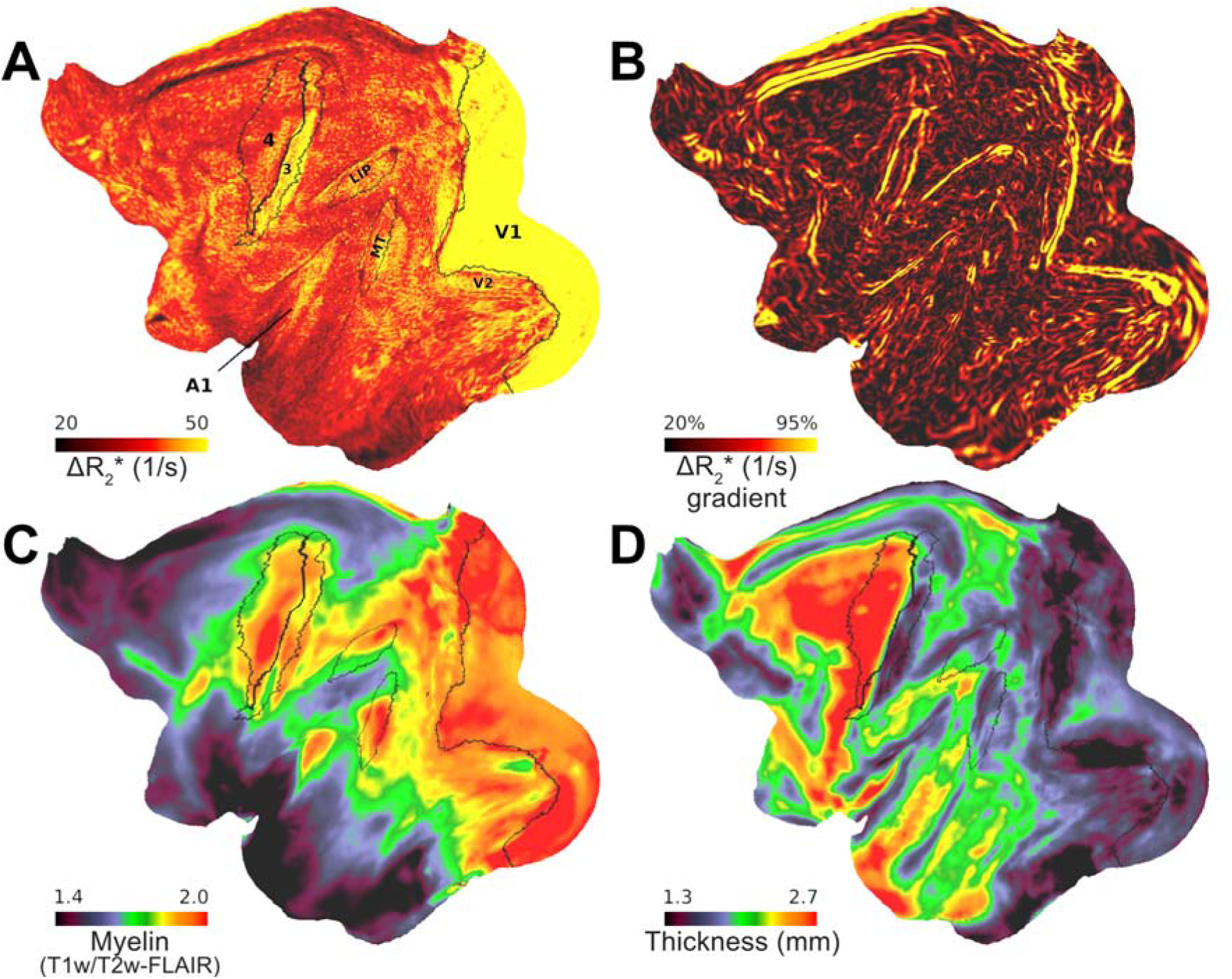
Variations in vascular network architecture reveal cortical area boundaries. **(A)** Ferumoxytol induced change in transverse relaxation rate (ΔR_2_*) displayed at a representative equivolumetric layer 4b (EL4b) (N=4). Overlaid black lines show exemplary M132 atlas area boundaries. **(B)** ΔR_2_* gradients co-align with exemplary areal boundaries. Red arrow indicates an artifact from inferior sagittal sinus. Average **(C)** mid-thickness weighted T1w/T2w-FLAIR myelin and **(D)** cortical thickness maps.

**Figure 5.**
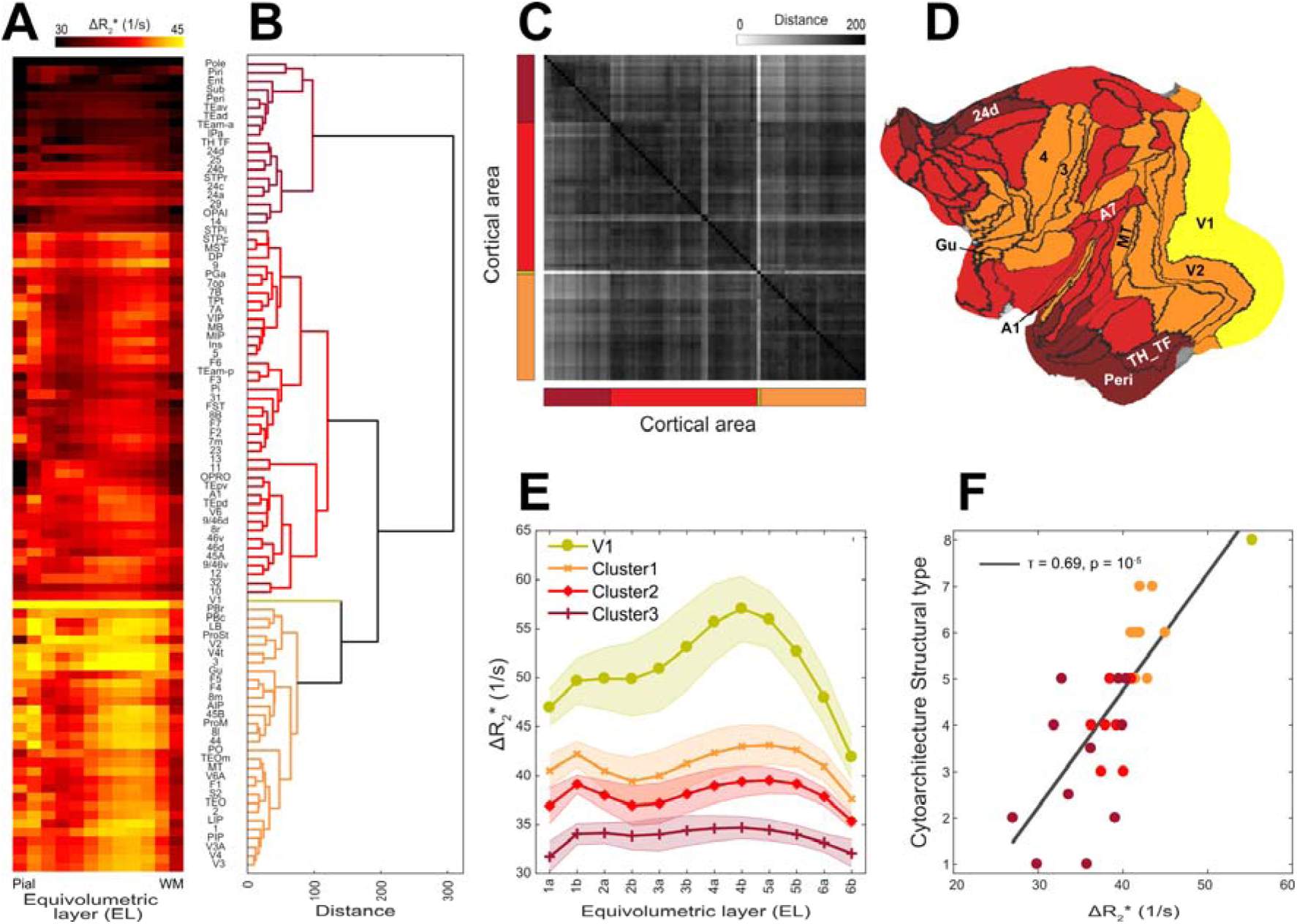
Hierarchical organization and principal types of cerebral vasculature. **(A)** Average ΔR_2_* equivolumetric layers (ELs) ascending from pial surface (left) to white matter surface (right) (N=4; hemispheres=8). Parcel order was sorted by **(B)** dendrogram determined using Wards’ method. **(C)** Similarity matrix as estimated using Euclidean distance. **(D)** Clusters displayed on a cortical flat-map. **(E)** Average cluster profiles. Error-bar indicates standard deviation across parcels within each cluster. **(F)** Cytoarchitectonic structural type co-vary with ΔR_2_*.

Because terminals of myelinated axons often overlap with high oxidative metabolism (Horton, 1984; Rockoff et al., 2014), we also examined the association between ΔR_2_* and T1w/T2w-FLAIR, an indirect proxy measure of cortical myelin density (Fig. 4A, C) (Glasser and Van Essen 2011, Autio et al., 2024). We found that M132 atlas parcellated ΔR_2_* was positively correlated with intra-cortical T1w/T2w-FLAIR myelin (R = 0.49 ± 0.16), and negatively correlated with cortical thickness (R = -0.37 ± 0.10) (Fig. 4D).

### 3.3 Translaminar vascular volume variations link with neuroanatomical organization

Across the cortical areas and layers, average ΔR_2_* profiles exhibited moderate variability (Fig. 5A, Supp. Fig. 5). To search for repetitive patterns in translaminar vascularity, we applied agglomerative clustering to concatenated group data. This analysis revealed distinct groups of vascularity arranged between eulaminate and agranular regions (Fig. 5B, C, D). A unique vascular profile was identified in V1, characterized by very dense vascularity and prominent peak density in EL4b (Fig. 5A, E). Average cluster profiles demonstrate that translaminar ΔR_2_* was relatively high in isocortical areas, and low in agranular areas (Fig. 5E). The cluster boundaries (Fig. 5D) typically occurred in vicinity of the strong ΔR_2_* gradients (Fig. 4B). Given the clustering between agranular and granular eulaminate cortices, we further corroborated whether the heterogeneous vascularization is associated with local microcircuit specialization of the cortex. For this objective, we used the designation of the cytoarchitectonic classification mapped onto M132 atlas (Burt et al., 2018; Hilgetag et al., 2016). This analysis confirmed that CBV, indeed, varies along the cytoarchitectonic types (Kendall’s tau τ = 0.69, *p* < 10^-5^) (Fig. 5F). We also found a close association between baseline R_2_* and cytoarchitecture (τ = 0.73, *p* < 10^-6^). In the isocortex, the majority of the areas exhibited a distinctively high ΔR_2_* in EL4 (Fig. 5A, E). The primary input layer, approximated as EL4a/b, exhibited systematically higher vascularity than in the primary output layer (*p* < 10^-25^), approximated as EL5a/b. In contrast, the majority of the agranular and dysgranular areas (cluster3; Fig. 5) exhibited weak laminar differentiation and a modest vascular density peak in EL1-2 (Fig. 5A).

Given neurons and receptors collectively constitute approximately 75-80% of the brain’s total energy budget (Howarth et al., 2012; Hyder et al., 2013), we next asked whether the regional variation in cerebrovascular network architecture (Fig. 1E) is associated with heterogeneous cellular and receptor densities. To address this question, we applied linear regression model, utilizing quantitative neuron (Collins et al., 2010) and receptor density maps (Froudist-Walsh, et al., 2023), to predict variation in ΔR_2_* (Fig. 6A, B, D). This analysis revealed a positive correlation between CBV and neuron density in the middle cortical layers, where neuron density is typically highest, while revealing a negative correlation with receptor density in the superficial layers where synaptic density is highest (Fig. 6F). Additionally, we observed that baseline R_2_* exhibited positive correlation with neuron density and negative correlation with receptor density (Fig. 6E).

**Figure 6.**
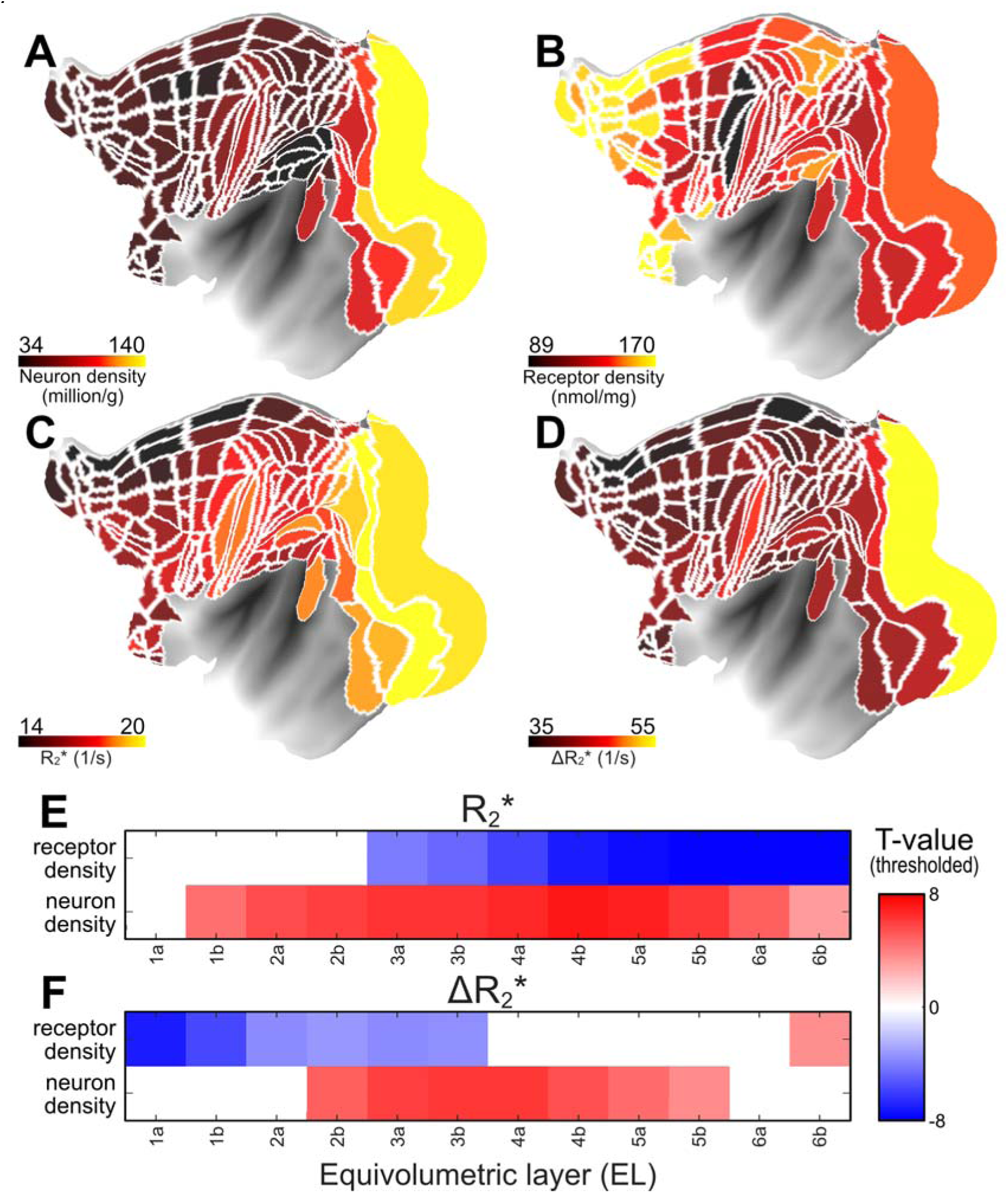
The anatomical underpinnings of the vascular network architecture. **(A)** Neuron (Collins et al., 2010), **(B)** total receptor density (Froudist-Walsh et al., 2023), **(C, D)** R_2_* and ΔR_2_* (current study). Multiple linear regression model was used to investigate the relationship between neuron and total receptor densities and **(E)** baseline R_2_* and **(F)** ΔR_2_* across layers. T-values are threshold at significance level (*p* < 0.05, Bonferroni corrected).

Because the Julich cortical area atlas covers only a section of the cerebral cortex, and the neuron density estimates are interpolated maps, we extended our analysis using the original Collins sample borders encompassing the entire cerebral cortex (Supp. Fig. 6A-C). This analysis reaffirmed the positive correlation with ΔR_2_* (peak at EL2, R = 0.80, *p* < 10^-11^) and baseline R_2_* (peak at EL2a, R = 0.86, *p* < 10^-13^), yielding linear coefficients of ΔR_2_* = 102 × 10^3^ neurons/s and R_2_* = 41 × 10^3^ neurons/s (Supp. Fig. 6D-G). This suggests that the sensitivity of quantitative layer R_2_* MRI in detecting neuronal loss is relatively weak, and the introduction of the Ferumoxytol contrast agent has the potential to enhance this sensitivity by a factor of 2.5.

Having established that vascular volume is associated with fundamental building units of cortical microcircuitry (Fig. 5F, Fig. 6E, F), our subsequent inquiry aimed to explore connection with interneurons that govern the neuroenergetics of local neural networks (Buzsáki et al., 2007). By utilizing the interneuron densities mapped onto M132 atlas (Burt et al., 2018), we identified a positive correlation between ΔR_2_* and parvalbumin interneuron density (peak at EL5b, R = 0.72, *p* < 10^-6^; Bonferroni corrected). In contrast, ΔR_2_* showed negative correlation with the density of calretinin-expressing slow-spiking interneurons which preferentially target distal dendrites (peak EL1b, R = -0.61, *p* < 0.001; Bonferroni corrected). In conjunction, we found that ΔR_2_* was also negatively correlated with dendritic tree size (R = -0.46, *p* < 0.01) (Supp. Fig. 7A, D, E), the number of spines in L3 pyramidal cells (R = -0.37, *p* = 0.06), and also with R_2_* (R = -0.69, *p* < 0.001) (Supp. Fig. 7B, D, F) (Elston 2002, Froudist-Walsh et al., 2023). These findings establish the intricate relationship between vascular density and the regulatory mechanisms governing diverse neural circuitry within the cerebral cortex.

## 4 Discussion

We present a noninvasive methodology to evaluate layer variations in vascular network architecture in the primate cerebral cortex. The quantitative cortical layer thickness adjusted ferumoxytol-weighted MRI enables exploration of systematic variations in cortical energy supply architecture and vessel-frequency informed image acquisition enables benchmarking penetrating vessel density measures relative to the anatomical ‘ground-truth’. These advances enabled us to unravel the systematic relation between vascularity and neurometabolic factors such as neuron and synaptic densities. Altogether, our study provides methodological and conceptual advancements in the field of cerebrovascular imaging.(Raichle and Mintun, 2006)

### 4.1 Methodological considerations - vessel density informed MRI

To gain insights into the organization of cerebrovascular networks, it is important to critically sample the large irrigating arteries and draining veins while preserving adequate SNR in gray matter. While the pial vessels can be directly visualized using high-resolution time-of-flight MRI (Bollmann et al., 2022), and computed tomography (Starosolski et al., 2015), imaging of the dense vascularity within the large and highly convoluted primate gray matter presents other formidable challenges. Here, we used a combination of ferumoxytol contrast agent and laminar-resolution 3D GRE MRI to map cerebrovascular architecture in macaque monkeys. These methods allowed us to indirectly delineate large vessels and estimate translaminar variations in cortical microvasculature.

This methodology, however, has known limitations. First, gradient-echo imaging is more sensitized toward large pial vessels running along the cortical surface and large penetrating vessels, which could differentially bias the estimation of ΔR_2_* across cortical layers (Fig. 2A, 2B) (Boxerman et al., 1995; Zhao et al., 2006). Additionally, vessel orientation relative to the B_0_ direction introduce strong layer-specific biases in quantitative ΔR_2_* measurements (Supp. Fig. 1C) (Lauwers et al., 2008; Ogawa et al., 1993; Viessmann et al., 2019). To address these concerns, we conducted necessary corrections for B_0_-orientation, obtained parcel median values and regressed linear-trend thereby mitigating the effect of undersampling large-caliber vessels across ELs (Fig. 2C, Supp. Fig. 1). These analytical solutions yielded ΔR_2_* V1 translaminar profiles that more closely resembled capillary rather than large-vessel volume profiles thus substantiating the validity of our methodology (Fig. 3F; Weber et al., 2008).

Ferumoxytol-weighted MRI of macaque cerebral cortex also enables the benchmarking of the methodological strengths and limitations to noninvasively measure vessel and vessel network length densities relative to the ‘ground-truth’. In macaque V1, large vessel length density (threshold at 8 μm diameter) is 138 mm/mm^3^ and mean area irrigated or drained by the vessels are about 0.26 and 0.4 mm^2^ for arteries and veins, respectively (Weber et al., 2008). Combined, the total vessel density (= artery/0.26 + vein/0.40 mm^2^) is 6.3 vessels/mm^2^ (Zheng et al., 1991; Weber et al., 2008), but see (Adams et al., 2015; Keller et al., 2011). Based on the former literature estimates, we hypothesized that isotropic voxel of 0.23 mm (19 voxels/mm^2^) may enable critical sampling of large vessels in accordance with sampling theorem (the sampling frequency equals to or is greater than twice the spatial frequency of the underlying anatomical detail in the image). In V1, we found an average vessel density of 2.2 ± 0.7 vessels/mm^2^ (Fig. 2E) which corresponds to ≈30% of the ‘ground-truth’ estimate (Weber et al., 2008). Using cortical thickness as a reference, we estimate that the vessel length density is ≈8 mm/mm^3^ which corresponds to a modest ≈10% of the ‘ground-truth’ (Weber et al., 2008). The latter underestimate may be attributed to under-sampling of the branching arteriole and venule networks. Indeed, anatomical studies accounting for branching patterns have reported much higher vessel densities up to 30 vessels/mm^2^ (Keller et al., 2011; Adams et al., 2015). Further investigations are warranted, taking into account critical sampling frequencies associated with vessel branching patterns (Duverney 1981), achieving higher SNR through ultra-high B_0_ MRI (Bolan et al., 2006; Harel et al., 2010; Kim et al., 2013) and utilize high-resolution single-plane sequences and prospective motion correction schemes to accurately characterize regional vessel densities. Such advancements hold promise for improving vessel quantification, classifications for veins and arteries and constructing detailed cortical surface maps of the vascular networks which may have diagnostic and neurosurgical utilities (Fig. 2A, B) (Iadecola, 2013; Qi and Roper, 2021; Sweeney et al., 2018).

### 4.2 Sharp transitions in microvasculature are indicative of cortical area- and layer-specific energy requirements

These methodological advances enabled us to unveil variations in vascular density within the primate cerebral cortex. Primary sensory cortices, known for their high energy demands (Kennedy et al., 1978), exhibit distinctive vascularization patterns (Fig. 1C, D, Fig. 3B). Notably, V1, area 3, auditory cortex, and also MT, all demonstrate elevated levels of CO enzymatic activity compared to surrounding cortical regions (Hackett et al., 1998; Horton, 1984; Huntley and Jones, 1991; Krubitzer et al., 2004; Matelli et al., 1985; Morel et al., 1993; Sincich et al., 2003). CO staining often reveals sharp transitions historically employed to delineate cortical area boundaries and modular features of the cortex. Our results corroborate the over three-decade-old, yet previously untested, hypothesis that some inter-areal boundaries may be determined by their microcapillary density using contrast-agent-weighted MRI (Zheng et al., 1991). Dense vascularity in these areas, the sharp gradient-ridges observed between surrounding areas and co-alignment with existing areal atlases further support this hypothesis (Fig. 4A, B).

Beyond the primary sensory areas, our observations extend to various smaller layer-specific vascular transitions (Fig. 5A, B). Specifically, the lateral intraparietal (LIP) area exhibits high CBV and gradient-ridges relative to ventral intraparietal (VIP) area and associative Brodmann area 7 (BA7). In EL5b, a strong gradient-ridge was observed distinguishing areas F4 and 44 from 45B and 8L/m. We also note weaker vascular transitions between areas such as 3a vs 3b, 5 vs anterior intraparietal (AIP) area, supplementary motor cortex (SII) vs insula, A1 vs medial belt and 46v vs 46d. However, our ability to confidently determine these borders is constrained by the presence of large vessels, as well as potential surface placement errors, and validating these areal boundaries would benefit from utilizing multimodal approaches.

In their work, Zheng and colleagues also proposed that modular features of the cortex, characterized by greater vascular density in the V1 CO blobs (42%) than interblobs, could be delineated using contrast-agent-weighted MRI (Zheng et al., 1991). Such a large vascular density difference should be well-within our contrast-to-noise and spatial sampling limitations. However, our results do not support this hypothesis, as we do not observe a distinct vascular peak at the spatial frequency of CO blobs (≈2.2 1/mm; Fig. 2D). Our results align with anatomical studies challenging the existence of high capillary density in the V1 blobs (Kernel et al., 2011, Adams et al., 2015).

The variation in vascular volume also has implications for statistical power in fMRI. For example, the pallidum exhibits lowest vascular volume (Fig. 1C) while in V1 EL4, the vascular volume is approximately twice as high (Fig. 3D). According to the classical single-tissue compartment model, CNR is optimized when TE is matched with T_2_*. Consequently, there is no single optimal TE for cerebral blood volume weighted fMRI. Multi-echo EPI acquisition (Kuroiwa et al., 2014; Poser and Norris, 2009) may provide a more balanced comparison for statistical power across different brain regions and cortical layers.

### 4.3 The vascular network architecture is intricately connected to the neuroanatomical organization within cerebral cortex

Given the fivefold variability in neuron density across the cortex (Cohen et al., 2010), one might expect that there is a corresponding variation in cerebral blood flow (CBF) and CBV (Fig. 6) (Tsai et al., 2007). In the cerebral cortex, neurons account for a significant portion (≈80-90%) of energy demand, with most of this energy allocated to signaling (≈80%) and maintaining membrane resting potentials (≈20%) (Attwell and Laughlin, 2001; Howarth et al., 2012). Since firing frequency is modulatory and the neural networks utilize distributed coding, the maintenance of resting-state membrane potential determines the minimal energy budget and the lower-limit for cerebral perfusion. Based on neuronal variability and energy dedicated to maintaining surface potential, this suggest an approximate (4 × 20% ≈) 80% variation in CBF and a resultant 25% variation in CBV across the cortex, in line with Grubbs’ law (CBV = 0.80 × CBF^0.38^) (Grubb et al., 1974). In the cerebellar cortex, neuron density is higher, and the resting potentials are thought to account for more than 50% of energy usage (Howarth et al., 2012), aligning with its higher vascular volume compared to the cerebral cortex (Fig. 1F). However, this is a simplified estimation, and a more comprehensive assessment would need to account for an aggregate of biophysical factors such as neuron types, neuron membrane surface area, firing rates, dendritic and synaptic densities (Fig. 6F-G), neurotransmitter recycling, and other cell types (Kageyama 1982; Elston and Rose 1997; Perge et al., 2009; Harris et al., 2012). Indeed, the majority of the mitochondria reside in the dendrites and synaptic transmission is widely acknowledged to drive the majority of the energy consumption and blood flow (Wong-Riley, 1989; Attwell et al., 2001).

Extrapolating cortical ΔR_2_* to zero neuron density results in a large intercept (∼35 1/s), corresponding to 60% of the maximum cortical CBV (57 1/s; Supp. Fig. 6F). This supports the view that most of the energy consumption occurs in the neuropil—comprising dendrites, synapses, and axons—which accounts for ∼80–90% of cortical gray matter volume, whereas neuronal somata constitute only ∼10–20% (Wong-Riley, 1989). Although neuronal cell bodies exhibit higher CO activity per unit volume due to their dense mitochondrial content, these results suggest their overall contribution to the total CBV per mm³ tissue remains lower than that of the neuropil, given the latter’s substantially larger volume fraction in cortical tissue.

Contrary to our initial expectations, we observed a relatively smaller CBV in regions and layers with high receptor density (Fig. 6B, D, F). This relationship extends to other factors, such as number of spines (putative excitatory inputs) and dendrite tree size across the entire cerebral cortex (Supp. Fig. 7) (Froudist-Walsh et al., 2023, Elston 2007). These results align with the work of Weber and colleagues, who reported a similar negative correlation between vascular length density and synaptic density, as well as a positive correlation with neuron density in macaque V1 across cortical layers (Weber et al., 2008). This relation is also compatible with the opposing relation between CBV and GABAergic (GABA, γ-aminobutyric acid) interneuron subtypes (Fig. 7G): parvalbumin-expressing fast-firing interneurons target perisomatic parts of pyramidal neurons, whereas calretinin-expressing slow-firing interneurons target distal-dendrites. The interneurons are also well-positioned to play an important role in integrating activity of large numbers of excitatory principal cells and translating this into neurovascular regulation of local microcirculation via subcortical pathways (Cauli et al., 2004). Another perspective on our results considers that a relative smaller faction of synapses may be simultaneously active in larger neurons, and neurons with large dendrites may exhibit lower excitability, supporting pattern separation necessary for higher-level functions (Chavlis et al., 2017; Hawkins and Ahmad, 2016). To comprehensively understand the factors contributing to the vascular organization of the brain, experimental disentanglement through multivariate analysis of laminar cell types and receptor densities is needed (Hayashi et al., 2021, Froudist-Walsh et al., 2023). Moreover, employing more advanced statistical modeling, including considerations for synapse-neuron interactions, may be important for refined evaluations.

Another key finding of this study was the strong correlation between baseline R_2_* and neuron density (Supp. Fig. 6D, E). While R_2_* is well known to be influenced by iron, myelin, and deoxyhemoglobin densities, this correlation peaks in the superficial layers (Supp. Fig. 6E), suggesting a link to CO activity and the accumulation of deoxygenated venous blood draining from all cortical layers toward the pial network. Notably, the absolute range of superficial R_2_* values (max - min ≈ 6 s⁻¹; Supp. Fig. 6D) is approximately 12-30 times larger than the ΔR_2_* observed during task-based BOLD fMRI at 3T (0.2-0.5 1/s) (Yablonskiy and Haacke, 1994). Since venous oxygenation is around 60% and task-induced changes in blood flow account for only 5%–10% of the brain’s resting blood flow (Raichle and Mintun, 2006), these results suggest that superficial R_2_* (Fig. 1D) may serve as a more accurate proxy for total deoxyhemoglobin content (and thus total oxygen consumption), which scales with the neuron density of the underlying cortical gray matter. Importantly, superficial layers may also provide a more specific measure of deoxyhemoglobin, as they are less influenced by myelin and iron, which are more concentrated in deeper cortical layers. Additionally, smaller but direct contributors, such as mitochondrial CO density—an iron-dependent factor—may also play a role in this relationship.

Additionally, our investigation also uncovered an overlap between vascular volume and myelin (Fig. 3A, C). Myelin may indirectly contribute to increased energy consumption in the cortical gray matter by facilitating higher frequency firing in comparison to unmyelinated axons (Perge et al., 2012; Saab et al., 2016). A large fraction of cortical myelin enwraps the axons of parvalbumin-expressing fast-spiking interneurons (Stedehouder et al., 2017). Although interneurons represent a minority of the neuronal population (10-15%), the parvalbumin-expressing interneurons’ ability to sustain high-frequency gamma oscillations (30-100 Hz) may match the sparse firing of the majority principal cells (Buzsáki et al., 2007). Indeed, parvalbumin-expressing interneurons exhibit higher mitochondrial volume compared to other cells in the brain and a 3-fold higher CO activity than principal neurons (Gulyás et al., 2006; Kageyama and Wong-Riley, 1982; Kann et al., 2014; Nie and Wong-Riley, 1995). Thus, the high metabolic load of parvalbumin-expressing interneurons makes them potentially vulnerable to failures in the vascular network due to aging, Alzheimer’s disease as well as stroke (Kann, 2016).

Given the distinct pre- and postsynaptic metabolic requirements, heterogenous translaminar vascularization may also indicate distinct cortical layers each characterized by anatomically and physiologically distinct feedforward and feedback pathways (Bastos et al., 2015; Borowsky and Collins, 1989; Kageyama and Wong-Riley, 1982; Takahata, 2016). A prime example is V1 where the primary input layer 4 (L4) has more dense vascularization and 50% higher CO activity in comparison with primary output L5 (Fig. 3F) (Keller et al., 2011; Livingstone and Hubel, 1982). This elevated energy demand in L4 may arise, in part, from spontaneous supra-threshold gamma-frequency oscillations between the retina→lateral geniculate nucleus→L4 (Castelo-Branco et al., 1998) along with recurrent amplification of local and distant inputs (Douglas and Martin, 2007; Shu et al., 2003). When viewed in terms of information flow, CBV appear to decrease along the canonical circuit pathway (e.g., L4→L2/3→L5) in the primary visual cortex (Douglas and Martin, 2007) and as one ascends the hierarchy (e.g., V1→V2→V3&4→MT→7A) from primary sensory areas (Fig. 3F, Supp. Fig. 8) (Felleman and Van Essen, 1991; Nikola T. Markov et al., 2014; N. T. Markov et al., 2014). A similar pattern is observed in the auditory hierarchy, where the inferior colliculus, an early processing hub, exhibits the highest vascular volume, followed by a gradual reduction along cortical auditory ‘where’ and ‘what’ pathways (Fig. 1F, Fig. 3B). In the agranular and dysgranular regions, characterized by a lack of distinct L4, the translaminar CBV profiles did not exhibit a distinct peak at around EL4 nor signatures of canonical circuitry (Fig. 5). These results demonstrate a greater allocation of the energy budget to early stages of feedforward processing in primary cortical areas characterized by strong sensory inputs, as opposed to higher-level cortical areas characterized by high synaptic densities supporting cognitive and behavioral functions (Fig. 6B) (Yokoyama et al., 2021).

The anatomical uniformity of the primate neocortex is thought to reflect extensive replication of a few specialized microcircuits varying along the brain’s hierarchical organization (Douglas and Martin, 2007; Hilgetag et al., 2016; Markram et al., 2004). Analogous to cortical circuitry, our study reveals the large-scale replication of translaminar vascular network motifs in primates (Fig. 4B, C, D). Since the cerebrovascular system evolved to support the high energy demands of neural information processing, this raises questions about whether the large-scale replication of anatomical and vascular circuits is evolutionarily coupled (Carmeliet and Tessier-Lavigne, 2005). In mice, comprehensive analysis of the cerebrovascular system has revealed distinct translaminar types between sensory and motor-integrative areas (Kirst et al., 2020). In macaque, we found the strongest distinction between isocortex and allocortex and its adjacent regions (Fig. 6B). The species difference may reflect evolutionary expansion and emergence of new cortical layers. For instance, in mice the primary somatosensory cortex exhibits highest vascularization (Kirst et al., 2020) whereas our results show that in macaque the highest vascularization is in the V1 (Fig. 5A, E) (Duverney 1981). According to the theory that sensory systems, behavior, and habitat choice are all influenced by evolutionary processes (Endler, 1992; Ikeda et al., 2023), this may reflect an evolutionary adaptation to an environment in which the visual landscape is an ecologically more important sensory domain in primates.

## Author contributions

Conceptualization: JAA; Methodology: JAA; Software: JAA, IK; Formal analysis: JAA, IK, TI; Investigation: JAA, TO, YM, MO; Writing - original draft: JAA; Writing - review & editing: JAA, IK, MFG, DVCE, TH; Visualization: JAA, IK, TI; Resources: JAA, YU, TH; Project administration: TO; Funding acquisition: JAA, TH.

## Acknowledgments

The authors appreciate discussions and technical contributions from Akiko Uematsu, Timothy S. Coalson, Katsutoshi Murata, and Reiko Kobayashi. This research is partially supported by JSPS KAKENHI Grant Number (JP20K15945, JAA), by the program for Brain/MINDS and Brain/MINDS-beyond from Japan Agency for Medical Research and development, AMED (JP18dm037006, TH) and by NIH R01MH60974 (DCVE, MFG). The authors have no conflicts of interest to declare.

## References

Adams, D.L., Piserchia, V., Economides, J.R., Horton, J.C., 2015. Vascular Supply of the Cerebral Cortex is Specialized for Cell Layers but Not Columns. Cereb. Cortex 25, 3673–3681. 10.1093/cercor/bhu221

Autio, J.A., Glasser, M.F., Ose, T., Donahue, C.J., Bastiani, M., Ohno, M., Kawabata, Y., Urushibata, Y., Murata, K., Nishigori, K., Yamaguchi, M., Hori, Y., Yoshida, A., Go, Y., Coalson, T.S., Jbabdi, S., Sotiropoulos, S.N., Kennedy, H., Smith, S., Van Essen, D.C., Hayashi, T., 2020. Towards HCP-Style macaque connectomes: 24-Channel 3T multi-array coil, MRI sequences and preprocessing. NeuroImage 215, 116800. 10.1016/j.neuroimage.2020.116800

Autio, J.A., Uematsu, A., Ikeda, T., Ose, T., Hou, Y., Magrou, L., Kimura, I., Ohno, M., Murata, K., Coalson, T., Kennedy, H., Glasser, M.F., Essen, D.C.V., Hayashi, T., 2024. Charting Cortical-Layer Specific Area Boundaries Using Gibbs Ringing Attenuated T1w/T2w-FLAIR Myelin MRI. 10.1101/2024.09.27.615294

Autio, J.A., Zhu, Q., Li, X., Glasser, M.F., Schwiedrzik, C.M., Fair, D.A., Zimmermann, J., Yacoub, E., Menon, R.S., Van Essen, D.C., Hayashi, T., Russ, B., Vanduffel, W., 2021. Minimal specifications for non-human primate MRI: Challenges in standardizing and harmonizing data collection. NeuroImage 236, 118082. 10.1016/j.neuroimage.2021.118082

Avadiappan, S., Payabvash, S., Morrison, M.A., Jakary, A., Hess, C.P., Lupo, J.M., 2020. A Fully Automated Method for Segmenting Arteries and Quantifying Vessel Radii on Magnetic Resonance Angiography Images of Varying Projection Thickness. Front. Neurosci. 14, 537. 10.3389/fnins.2020.00537

Bastos, A.M., Vezoli, J., Bosman, C.A., Schoffelen, J.-M., Oostenveld, R., Dowdall, J.R., De Weerd, P., Kennedy, H., Fries, P., 2015. Visual Areas Exert Feedforward and Feedback Influences through Distinct Frequency Channels. Neuron 85, 390–401. 10.1016/j.neuron.2014.12.018

Bolan, P.J., Yacoub, E., Garwood, M., Ugurbil, K., Harel, N., 2006. In vivo micro-MRI of intracortical neurovasculature. NeuroImage 32, 62–69. 10.1016/j.neuroimage.2006.03.027

Bollmann, S., Mattern, H., Bernier, M., Robinson, S.D., Park, D., Speck, O., Polimeni, J.R., 2022. Imaging of the pial arterial vasculature of the human brain in vivo using high-resolution 7T time-of-flight angiography. eLife 11, e71186. 10.7554/eLife.71186

Borowsky, I.W., Collins, R.C., 1989. Metabolic anatomy of brain: A comparison of regional capillary density, glucose metabolism, and enzyme activities. J. Comp. Neurol. 288, 401–413. 10.1002/cne.902880304

Boxerman, J.L., Hamberg, L.M., Rosen, B.R., Weisskoff, R.M., 1995. Mr contrast due to intravascular magnetic susceptibility perturbations. Magn. Reson. Med. 34, 555–566. 10.1002/mrm.1910340412

Buch, S., Chen, Y., Jella, P., Ge, Y., Haacke, E.M., 2022. Vascular mapping of the human hippocampus using Ferumoxytol-enhanced MRI. NeuroImage 250, 118957. 10.1016/j.neuroimage.2022.118957

Burt, J.B., Demirtaş, M., Eckner, W.J., Navejar, N.M., Ji, J.L., Martin, W.J., Bernacchia, A., Anticevic, A., Murray, J.D., 2018. Hierarchy of transcriptomic specialization across human cortex captured by structural neuroimaging topography. Nat. Neurosci. 21, 1251–1259. 10.1038/s41593-018-0195-0

Buzsáki, G., Kaila, K., Raichle, M., 2007. Inhibition and Brain Work. Neuron 56, 771–783. 10.1016/j.neuron.2007.11.008

Carmeliet, P., Tessier-Lavigne, M., 2005. Common mechanisms of nerve and blood vessel wiring. Nature 436, 193–200. 10.1038/nature03875

Castelo-Branco, M., Neuenschwander, S., Singer, W., 1998. Synchronization of visual responses between the cortex, lateral geniculate nucleus, and retina in the anesthetized cat. J. Neurosci. Off. J. Soc. Neurosci. 18, 6395–6410. 10.1523/JNEUROSCI.18-16-06395.1998

Cauli, B., Tong, X.-K., Rancillac, A., Serluca, N., Lambolez, B., Rossier, J., Hamel, E., 2004. Cortical GABA Interneurons in Neurovascular Coupling: Relays for Subcortical Vasoactive Pathways. J. Neurosci. 24, 8940–8949. 10.1523/JNEUROSCI.3065-04.2004

Chavlis, S., Petrantonakis, P.C., Poirazi, P., 2017. Dendrites of dentate gyrus granule cells contribute to pattern separation by controlling sparsity. Hippocampus 27, 89–110. 10.1002/hipo.22675

Collins, C.E., Airey, D.C., Young, N.A., Leitch, D.B., Kaas, J.H., 2010. Neuron densities vary across and within cortical areas in primates. Proc. Natl. Acad. Sci. 107, 15927–15932. 10.1073/pnas.1010356107

Condé, F., Lund, J.S., Jacobowitz, D.M., Baimbridge, K.G., Lewis, D.A., 1994. Local circuit neurons immunoreactive for calretinin, calbindin D-28k or parvalbumin in monkey prefrontal cortex: distribution and morphology. J. Comp. Neurol. 341, 95–116. 10.1002/cne.903410109

Douglas, R.J., Martin, K.A.C., 2007. Recurrent neuronal circuits in the neocortex. Curr. Biol. 17, R496–R500. 10.1016/j.cub.2007.04.024

Duvernoy, H.M., Delon, S., Vannson, J.L., 1981. Cortical blood vessels of the human brain. Brain Res. Bull. 7, 519–579. 10.1016/0361-9230(81)90007-1

Elston, G.N., 2007. in Evolution of Nervous Systems (eds Kaas, J. H. & Preuss, T. M.). Elsevier, pp. 191–242.

Elston, G.N., 2002. Cortical heterogeneity: implications for visual processing and polysensory integration. J. Neurocytol. 31, 317–335. 10.1023/a:1024182228103

Endler, J.A., 1992. Signals, Signal Conditions, and the Direction of Evolution. Am. Nat. 139, S125–S153.

Felleman, D.J., Van Essen, D., 1991. Distributed hierarchical processing in the primate cerebral cortex. Cereb. Cortex N. Y. N 1991 1, 1–47. 10.1093/cercor/1.1.1

Frangi, A.F., Niessen, W.J., Vincken, K.L., Viergever, M.A., 1998. Multiscale vessel enhancement filtering, in: Wells, W.M., Colchester, A., Delp, S. (Eds.), Medical Image Computing and Computer-Assisted Intervention — MICCAI’98, Lecture Notes in Computer Science. Springer, Berlin, Heidelberg, pp. 130–137. 10.1007/BFb0056195

Froudist-Walsh, S., Bliss, D.P., Ding, X., Rapan, L., Niu, M., Knoblauch, K., Zilles, K., Kennedy, H., Palomero-Gallagher, N., Wang, X.-J., 2021. A dopamine gradient controls access to distributed working memory in the large-scale monkey cortex. Neuron 109, 3500–3520.e13. 10.1016/j.neuron.2021.08.024

Froudist-Walsh, S., Xu, T., Niu, M., Rapan, L., Zhao, L., Margulies, D.S., Zilles, K., Wang, X.-J., Palomero-Gallagher, N., 2023. Gradients of neurotransmitter receptor expression in the macaque cortex. Nat. Neurosci. 26, 1281–1294. 10.1038/s41593-023-01351-2

Gabbott, P.L., Bacon, S.J., 1996. Local circuit neurons in the medial prefrontal cortex (areas 24a,b,c, 25 and 32) in the monkey: II. Quantitative areal and laminar distributions. J. Comp. Neurol. 364, 609–636. 10.1002/(SICI)1096-9861(19960122)364:4<609::AID-CNE2>3.0.CO;2-7

Glasser, M.F., Sotiropoulos, S.N., Wilson, J.A., Coalson, T.S., Fischl, B., Andersson, J.L., Xu, J., Jbabdi, S., Webster, M., Polimeni, J.R., Van Essen, D.C., Jenkinson, M., 2013. The minimal preprocessing pipelines for the Human Connectome Project. NeuroImage 80, 105–124. 10.1016/j.neuroimage.2013.04.127

Glasser, M.F., Van Essen, D.C., 2011. Mapping human cortical areas in vivo based on myelin content as revealed by T1- and T2-weighted MRI. J. Neurosci. Off. J. Soc. Neurosci. 31, 11597–11616. 10.1523/JNEUROSCI.2180-11.2011

Greve, D.N., Fischl, B., 2009. Accurate and robust brain image alignment using boundary-based registration. NeuroImage 48, 63–72. 10.1016/j.neuroimage.2009.06.060

Gulyás, A.I., Buzsáki, G., Freund, T.F., Hirase, H., 2006. Populations of hippocampal inhibitory neurons express different levels of cytochrome c. Eur. J. Neurosci. 23, 2581–2594. 10.1111/j.1460-9568.2006.04814.x

Hackett, T. a., Stepniewska, I., Kaas, J. h., 1998. Subdivisions of auditory cortex and ipsilateral cortical connections of the parabelt auditory cortex in macaque monkeys. J. Comp. Neurol. 394, 475–495. 10.1002/(SICI)1096-9861(19980518)394:4<475::AID-CNE6>3.0.CO;2-Z

Harel, N., Bolan, P.J., Turner, R., Ugurbil, K., Yacoub, E., 2010. Recent Advances in High-Resolution MR Application and Its Implications for Neurovascular Coupling Research. Front. Neuroenergetics 2, 130. 10.3389/fnene.2010.00130

Harrison, R.V., Harel, N., Panesar, J., Mount, R.J., 2002. Blood capillary distribution correlates with hemodynamic-based functional imaging in cerebral cortex. Cereb. Cortex N. Y. N 1991 12, 225–233. 10.1093/cercor/12.3.225

Hawkins, J., Ahmad, S., 2016. Why Neurons Have Thousands of Synapses, a Theory of Sequence Memory in Neocortex. Front. Neural Circuits 10, 23. 10.3389/fncir.2016.00023

Hayashi, T., Hou, Y., Glasser, M.F., Autio, J.A., Knoblauch, K., Inoue-Murayama, M., Coalson, T., Yacoub, E., Smith, S., Kennedy, H., Van Essen, D.C., 2021. The nonhuman primate neuroimaging and neuroanatomy project. NeuroImage 229, 117726. 10.1016/j.neuroimage.2021.117726

Hilgetag, C.C., Medalla, M., Beul, S.F., Barbas, H., 2016. The primate connectome in context: Principles of connections of the cortical visual system. NeuroImage 134, 685–702. 10.1016/j.neuroimage.2016.04.017

Horton, J.C., 1984. Cytochrome oxidase patches: a new cytoarchitectonic feature of monkey visual cortex. Philos. Trans. R. Soc. Lond. B Biol. Sci. 10.1098/rstb.1984.0021

Howarth, C., Gleeson, P., Attwell, D., 2012. Updated energy budgets for neural computation in the neocortex and cerebellum. J. Cereb. Blood Flow Metab. 32, 1222–1232. 10.1038/jcbfm.2012.35

Huntley, G.W., Jones, E.G., 1991. The emergence of architectonic field structure and areal borders in developing monkey sensorimotor cortex. Neuroscience 44, 287–310. 10.1016/0306-4522(91)90055-s

Hyder, F., Rothman, D.L., Bennett, M.R., 2013. Cortical energy demands of signaling and nonsignaling components in brain are conserved across mammalian species and activity levels. Proc. Natl. Acad. Sci. U. S. A. 110, 3549–3554. 10.1073/pnas.1214912110

Iadecola, C., 2013. The pathobiology of vascular dementia. Neuron 80, 10.1016/j.neuron.2013.10.008.

Ikeda, T., Autio, J.A., Kawasaki, A., Takeda, C., Ose, T., Takada, M., Van Essen, D.C., Glasser, M.F., Hayashi, T., 2023. Cortical adaptation of the night monkey to a nocturnal niche environment: a comparative non-invasive T1w/T2w myelin study. Brain Struct. Funct. 228, 1107–1123. 10.1007/s00429-022-02591-x

Jenkinson, M., Bannister, P., Brady, M., Smith, S., 2002. Improved Optimization for the Robust and Accurate Linear Registration and Motion Correction of Brain Images. NeuroImage 17, 825–841. 10.1006/nimg.2002.1132

Ji, X., Ferreira, T., Friedman, B., Liu, R., Liechty, H., Bas, E., Chandrashekar, J., Kleinfeld, D., 2021. Brain microvasculature has a common topology with local differences in geometry that match metabolic load. Neuron 109, 1168–1187.e13. 10.1016/j.neuron.2021.02.006

Kageyama, G.H., Wong-Riley, M.T.T., 1982. Histochemical localization of cytochrome oxidase in the hippocampus: Correlation with specific neuronal types and afferent pathways. Neuroscience 7, 2337–2361. 10.1016/0306-4522(82)90199-3

Kann, O., 2016. The interneuron energy hypothesis: Implications for brain disease. Neurobiol. Dis. 90, 75–85. 10.1016/j.nbd.2015.08.005

Kann, O., Papageorgiou, I.E., Draguhn, A., 2014. Highly energized inhibitory interneurons are a central element for information processing in cortical networks. J. Cereb. Blood Flow Metab. Off. J. Int. Soc. Cereb. Blood Flow Metab. 34, 1270–1282. 10.1038/jcbfm.2014.104

Keller, A.L., Schüz, A., Logothetis, N.K., Weber, B., 2011. Vascularization of cytochrome oxidase-rich blobs in the primary visual cortex of squirrel and macaque monkeys. J. Neurosci. Off. J. Soc. Neurosci. 31, 1246–1253. 10.1523/JNEUROSCI.2765-10.2011

Kim, S.-G., Harel, N., Jin, T., Kim, T., Lee, P., Zhao, F., 2013. Cerebral blood volume MRI with intravascular superparamagnetic iron oxide nanoparticles. NMR Biomed. 26, 949–962. 10.1002/nbm.2885

Kirst, C., Skriabine, S., Vieites-Prado, A., Topilko, T., Bertin, P., Gerschenfeld, G., Verny, F., Topilko, P., Michalski, N., Tessier-Lavigne, M., Renier, N., 2020. Mapping the Fine-Scale Organization and Plasticity of the Brain Vasculature. Cell 180, 780–795.e25. 10.1016/j.cell.2020.01.028

Kondo, H., Tanaka, K., Hashikawa, T., Jones, E.G., 1999. Neurochemical gradients along monkey sensory cortical pathways: calbindin-immunoreactive pyramidal neurons in layers II and III. Eur. J. Neurosci. 11, 4197–4203. 10.1046/j.1460-9568.1999.00844.x

Krubitzer, L., Huffman, K.J., Disbrow, E., Recanzone, G., 2004. Organization of area 3a in macaque monkeys: Contributions to the cortical phenotype. J. Comp. Neurol. 471, 97–111. 10.1002/cne.20025

Kuroiwa, D., Obata, T., Kawaguchi, H., Autio, J., Hirano, M., Aoki, I., Kanno, I., Kershaw, J., 2014. Signal contributions to heavily diffusion-weighted functional magnetic resonance imaging investigated with multi-SE-EPI acquisitions. NeuroImage 98, 258–265. 10.1016/j.neuroimage.2014.04.050

Langkammer, C., Schweser, F., Krebs, N., Deistung, A., Goessler, W., Scheurer, E., Sommer, K., Reishofer, G., Yen, K., Fazekas, F., Ropele, S., Reichenbach, J.R., 2012. Quantitative susceptibility mapping (QSM) as a means to measure brain iron? A post mortem validation study. Neuroimage 62, 1593–1599. 10.1016/j.neuroimage.2012.05.049

Lauwers, F., Cassot, F., Lauwers-Cances, V., Puwanarajah, P., Duvernoy, H., 2008. Morphometry of the human cerebral cortex microcirculation: General characteristics and space-related profiles. NeuroImage 39, 936–948. 10.1016/j.neuroimage.2007.09.024

Lee, J., van Gelderen, P., Kuo, L.-W., Merkle, H., Silva, A.C., Duyn, J.H., 2011. T2*-based fiber orientation mapping. NeuroImage 57, 225–234. 10.1016/j.neuroimage.2011.04.026

Lewis, J.W., Essen, D.C.V., 2000. Mapping of architectonic subdivisions in the macaque monkey, with emphasis on parieto-occipital cortex. J. Comp. Neurol. 428, 79–111. 10.1002/1096-9861(20001204)428:1<79::AID-CNE7>3.0.CO;2-Q

Livingstone, M.S., Hubel, D.H., 1982. Thalamic inputs to cytochrome oxidase-rich regions in monkey visual cortex. Proc. Natl. Acad. Sci. U. S. A. 79, 6098–6101. 10.1073/pnas.79.19.6098

Markov, N. T., Ercsey-Ravasz, M.M., Ribeiro Gomes, A.R., Lamy, C., Magrou, L., Vezoli, J., Misery, P., Falchier, A., Quilodran, R., Gariel, M.A., Sallet, J., Gamanut, R., Huissoud, C., Clavagnier, S., Giroud, P., Sappey-Marinier, D., Barone, P., Dehay, C., Toroczkai, Z., Knoblauch, K., Van Essen, D.C., Kennedy, H., 2014. A Weighted and Directed Interareal Connectivity Matrix for Macaque Cerebral Cortex. Cereb. Cortex 24, 17–36. 10.1093/cercor/bhs270

Markov, Nikola T., Vezoli, J., Chameau, P., Falchier, A., Quilodran, R., Huissoud, C., Lamy, C., Misery, P., Giroud, P., Ullman, S., Barone, P., Dehay, C., Knoblauch, K., Kennedy, H., 2014. Anatomy of hierarchy: feedforward and feedback pathways in macaque visual cortex. J. Comp. Neurol. 522, 225–259. 10.1002/cne.23458

Markram, H., Toledo-Rodriguez, M., Wang, Y., Gupta, A., Silberberg, G., Wu, C., 2004. Interneurons of the neocortical inhibitory system. Nat. Rev. Neurosci. 5, 793–807. 10.1038/nrn1519

Matelli, M., Luppino, G., Rizzolatti, G., 1985. Patterns of cytochrome oxidase activity in the frontal agranular cortex of the macaque monkey. Behav. Brain Res. 18, 125–136. 10.1016/0166-4328(85)90068-3

Morel, A., Garraghty, P.E., Kaas, J.H., 1993. Tonotopic organization, architectonic fields, and connections of auditory cortex in macaque monkeys. J. Comp. Neurol. 335, 437–459. 10.1002/cne.903350312

Muehe, A.M., Feng, D., von Eyben, R., Luna-Fineman, S., Link, M.P., Muthig, T., Huddleston, A.E., Neuwelt, E.A., Daldrup-Link, H.E., 2016. Safety Report of Ferumoxytol for Magnetic Resonance Imaging in Children and Young Adults. Invest. Radiol. 51, 221–227. 10.1097/RLI.0000000000000230

Nie, F., Wong-Riley, M.T., 1995. Double labeling of GABA and cytochrome oxidase in the macaque visual cortex: quantitative EM analysis. J. Comp. Neurol. 356, 115–131. 10.1002/cne.903560108

Ogawa, S., Menon, R.S., Tank, D.W., Kim, S.G., Merkle, H., Ellermann, J.M., Ugurbil, K., 1993. Functional brain mapping by blood oxygenation level-dependent contrast magnetic resonance imaging. A comparison of signal characteristics with a biophysical model. Biophys. J. 64, 803–812.

Perge, J.A., Niven, J.E., Mugnaini, E., Balasubramanian, V., Sterling, P., 2012. Why Do Axons Differ in Caliber? J. Neurosci. 32, 626–638. 10.1523/JNEUROSCI.4254-11.2012

Poser, B.A., Norris, D.G., 2009. Investigating the benefits of multi-echo EPI for fMRI at 7 T. NeuroImage 45, 1162–1172. 10.1016/j.neuroimage.2009.01.007

Qi, Y., Roper, M., 2021. Control of low flow regions in the cortical vasculature determines optimal arterio-venous ratios. Proc. Natl. Acad. Sci. 118, e2021840118. 10.1073/pnas.2021840118

Raichle, M.E., Mintun, M.A., 2006. BRAIN WORK AND BRAIN IMAGING. Annu. Rev. Neurosci. 29, 449–476. 10.1146/annurev.neuro.29.051605.112819

Reina-De La Torre, F., Rodriguez-Baeza, A., Sahuquillo-Barris, J., 1998. Morphological characteristics and distribution pattern of the arterial vessels in human cerebral cortex: A scanning electron microscope study. Anat. Rec. 251, 87–96. 10.1002/(SICI)1097-0185(199805)251:1<87::AID-AR14>3.0.CO;2-7

Robinson, E.C., Garcia, K., Glasser, M.F., Chen, Z., Coalson, T.S., Makropoulos, A., Bozek, J., Wright, R., Schuh, A., Webster, M., Hutter, J., Price, A., Cordero Grande, L., Hughes, E., Tusor, N., Bayly, P.V., Van Essen, D.C., Smith, S.M., Edwards, A.D., Hajnal, J., Jenkinson, M., Glocker, B., Rueckert, D., 2018. Multimodal surface matching with higher-order smoothness constraints. NeuroImage 167, 453–465. 10.1016/j.neuroimage.2017.10.037

Robinson, E.C., Jbabdi, S., Glasser, M.F., Andersson, J., Burgess, G.C., Harms, M.P., Smith, S.M., Essen, D.C.V., Jenkinson, M., 2014. MSM: a new flexible framework for Multimodal Surface Matching⋆. NeuroImage 100, 414. 10.1016/j.neuroimage.2014.05.069

Rockoff, E.C., Balaram, P., Kaas, J.H., 2014. Patchy distributions of myelin and vesicular glutamate transporter 2 align with cytochrome oxidase blobs and interblobs in the superficial layers of the primary visual cortex. Eye Brain 6, 19–27. 10.2147/EB.S59797

Saab, A.S., Tzvetavona, I.D., Trevisiol, A., Baltan, S., Dibaj, P., Kusch, K., Möbius, W., Goetze, B., Jahn, H.M., Huang, W., Steffens, H., Schomburg, E.D., Pérez-Samartín, A., Pérez-Cerdá, F., Bakhtiari, D., Matute, C., Löwel, S., Griesinger, C., Hirrlinger, J., Kirchhoff, F., Nave, K.-A., 2016. Oligodendroglial NMDA Receptors Regulate Glucose Import and Axonal Energy Metabolism. Neuron 91, 119–132. 10.1016/j.neuron.2016.05.016

Schmid, F., Barrett, M.J.P., Jenny, P., Weber, B., 2019. Vascular density and distribution in neocortex. NeuroImage 197, 792–805. 10.1016/j.neuroimage.2017.06.046

Shu, Y., Hasenstaub, A., McCormick, D.A., 2003. Turning on and off recurrent balanced cortical activity. Nature 423, 288–293. 10.1038/nature01616

Sincich, L.C., Adams, D.L., Horton, J.C., 2003. Complete flatmounting of the macaque cerebral cortex. Vis. Neurosci. 20, 663–686. 10.1017/s0952523803206088

Starosolski, Z., Villamizar, C.A., Rendon, D., Paldino, M.J., Milewicz, D.M., Ghaghada, K.B., Annapragada, A.V., 2015. Ultra High-Resolution In vivo Computed Tomography Imaging of Mouse Cerebrovasculature Using a Long Circulating Blood Pool Contrast Agent. Sci. Rep. 5, 10178. 10.1038/srep10178

Stedehouder, J., Couey, J.J., Brizee, D., Hosseini, B., Slotman, J.A., Dirven, C.M.F., Shpak, G., Houtsmuller, A.B., Kushner, S.A., 2017. Fast-spiking Parvalbumin Interneurons are Frequently Myelinated in the Cerebral Cortex of Mice and Humans. Cereb. Cortex N. Y. N 1991 27, 5001–5013. 10.1093/cercor/bhx203

Sweeney, M.D., Kisler, K., Montagne, A., Toga, A.W., Zlokovic, B.V., 2018. The role of brain vasculature in neurodegenerative disorders. Nat. Neurosci. 21, 1318–1331. 10.1038/s41593-018-0234-x

Tabelow, K., Balteau, E., Ashburner, J., Callaghan, M.F., Draganski, B., Helms, G., Kherif, F., Leutritz, T., Lutti, A., Phillips, C., Reimer, E., Ruthotto, L., Seif, M., Weiskopf, N., Ziegler, G., Mohammadi, S., 2019. hMRI - A toolbox for quantitative MRI in neuroscience and clinical research. NeuroImage 194, 191–210. 10.1016/j.neuroimage.2019.01.029

Takahata, T., 2016. What Does Cytochrome Oxidase Histochemistry Represent in the Visual Cortex? Front. Neuroanat. 10.

Toledo, J.B., Arnold, S.E., Raible, K., Brettschneider, J., Xie, S.X., Grossman, M., Monsell, S.E., Kukull, W.A., Trojanowski, J.Q., 2013. Contribution of cerebrovascular disease in autopsy confirmed neurodegenerative disease cases in the National Alzheimer’s Coordinating Centre. Brain 136, 2697–2706. 10.1093/brain/awt188

Tsai, P.S., Kaufhold, J.P., Blinder, P., Friedman, B., Drew, P.J., Karten, H.J., Lyden, P.D., Kleinfeld, D., 2009. Correlations of Neuronal and Microvascular Densities in Murine Cortex Revealed by Direct Counting and Colocalization of Nuclei and Vessels. J. Neurosci. 29, 14553–14570. 10.1523/JNEUROSCI.3287-09.2009

Van Essen, D.C., Maunsell, J.H.R., 1980. Two-dimensional maps of the cerebral cortex. J. Comp. Neurol. 191, 255–281. 10.1002/cne.901910208

Van Essen, D.C., Smith, J., Glasser, M.F., Elam, J., Donahue, C.J., Dierker, D.L., Reid, E.K., Coalson, T., Harwell, J., 2017. The Brain Analysis Library of Spatial maps and Atlases (BALSA) database. NeuroImage, Data Sharing Part II 144, 270–274. 10.1016/j.neuroimage.2016.04.002

Viessmann, O., Scheffler, K., Bianciardi, M., Wald, L.L., Polimeni, J.R., 2019. Dependence of resting-state fMRI fluctuation amplitudes on cerebral cortical orientation relative to the direction of B0 and anatomical axes. NeuroImage 196, 337–350. 10.1016/j.neuroimage.2019.04.036

Weber, B., Keller, A.L., Reichold, J., Logothetis, N.K., 2008. The Microvascular System of the Striate and Extrastriate Visual Cortex of the Macaque. Cereb. Cortex 18, 2318–2330. 10.1093/cercor/bhm259

Wong-Riley, M.T.T., 1989. Cytochrome oxidase: an endogenous metabolic marker for neuronal activity. Trends Neurosci. 12, 94–101. 10.1016/0166-2236(89)90165-3

Yablonskiy, D.A., Haacke, E.M., 1994. Theory of NMR signal behavior in magnetically inhomogeneous tissues: the static dephasing regime. Magn. Reson. Med. 32, 749–763. 10.1002/mrm.1910320610

Yokoyama, C., Autio, J.A., Ikeda, T., Sallet, J., Mars, R.B., Van Essen, D.C., Glasser, M.F., Sadato, N., Hayashi, T., 2021. Comparative connectomics of the primate social brain. NeuroImage 245, 118693. 10.1016/j.neuroimage.2021.118693

Zhang, Y., Brady, M., Smith, S., 2001. Segmentation of brain MR images through a hidden Markov random field model and the expectation-maximization algorithm. IEEE Trans. Med. Imaging 20, 45–57. 10.1109/42.906424

Zhao, F., Wang, P., Hendrich, K., Ugurbil, K., Kim, S.-G., 2006. Cortical layer-dependent BOLD and CBV responses measured by spin-echo and gradient-echo fMRI: Insights into hemodynamic regulation. NeuroImage 30, 1149–1160. 10.1016/j.neuroimage.2005.11.013

Zheng, D., LaMantia, A.S., Purves, D., 1991. Specialized vascularization of the primate visual cortex. J. Neurosci. Off. J. Soc. Neurosci. 11, 2622–2629.

