## Supplemental Figure 1-8 for "Mapping vascular network architecture in primate brain using ferumoxytol-weighted laminar MRI"

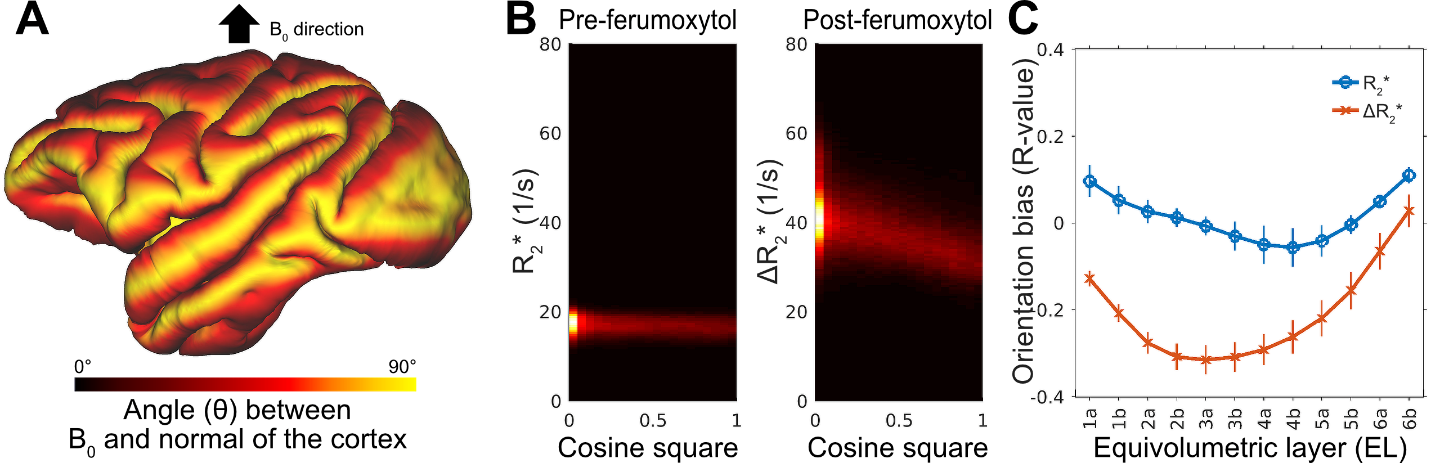

**Supplementary Figure 1. Transverse relaxation rate (R_2_*) measures are biased by the orientation of the static magnetic field (B_0_). (A)** The angle (θ) between B_0_ and the normal of the cortex. **(B)** Representative pre-ferumoxytol R_2_* (left) and ferumoxytol induced change in R_2_* (ΔR_2_*; right) plotted with respect to the cosine squared of θ from an equivolumetric layer 4a (EL4a) (N=1). Note that ΔR_2_* is high when the cortex and B_0_ are perpendicular and low when they are parallel. **(C)** B_0_ orientation bias exhibits laminar depth dependence. Values indicate mean and error-bars indicate standard deviation across subjects (N=4). Interestingly, R_2_* is positively whereas ΔR_2_* is negatively correlated with B_0_ orientation. R_2_* bias may reflect diamagnetic myelin sheath enwrapping axons that are oriented mainly parallel to the normal of cortex whereas ΔR_2_* may reflect vessels (e.g arterioles, capillaries and venules) with a net orientation perpendicular to the normal of the cortex.

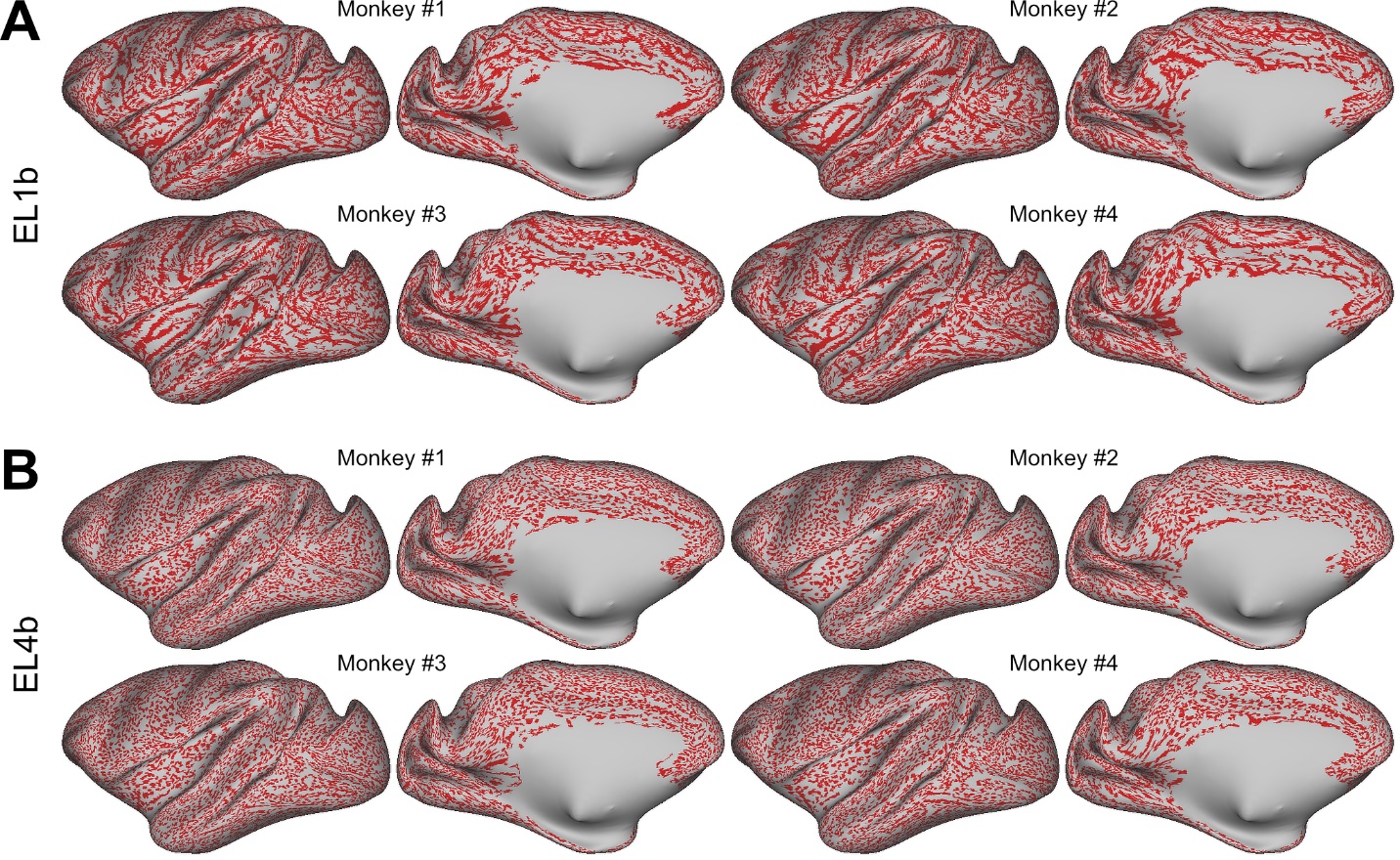

**Supplementary Figure 2.** **Consistency of pial vessel network mapping across subjects.** **(A)** After ferumoxytol contrast agent injection, pial vessels induce continuous signal-losses in the superficial equivolumetric layer 1b (EL1b). **(B)** Continuous signal-dropouts are largely absent in deeper cortical layers such as EL4b.

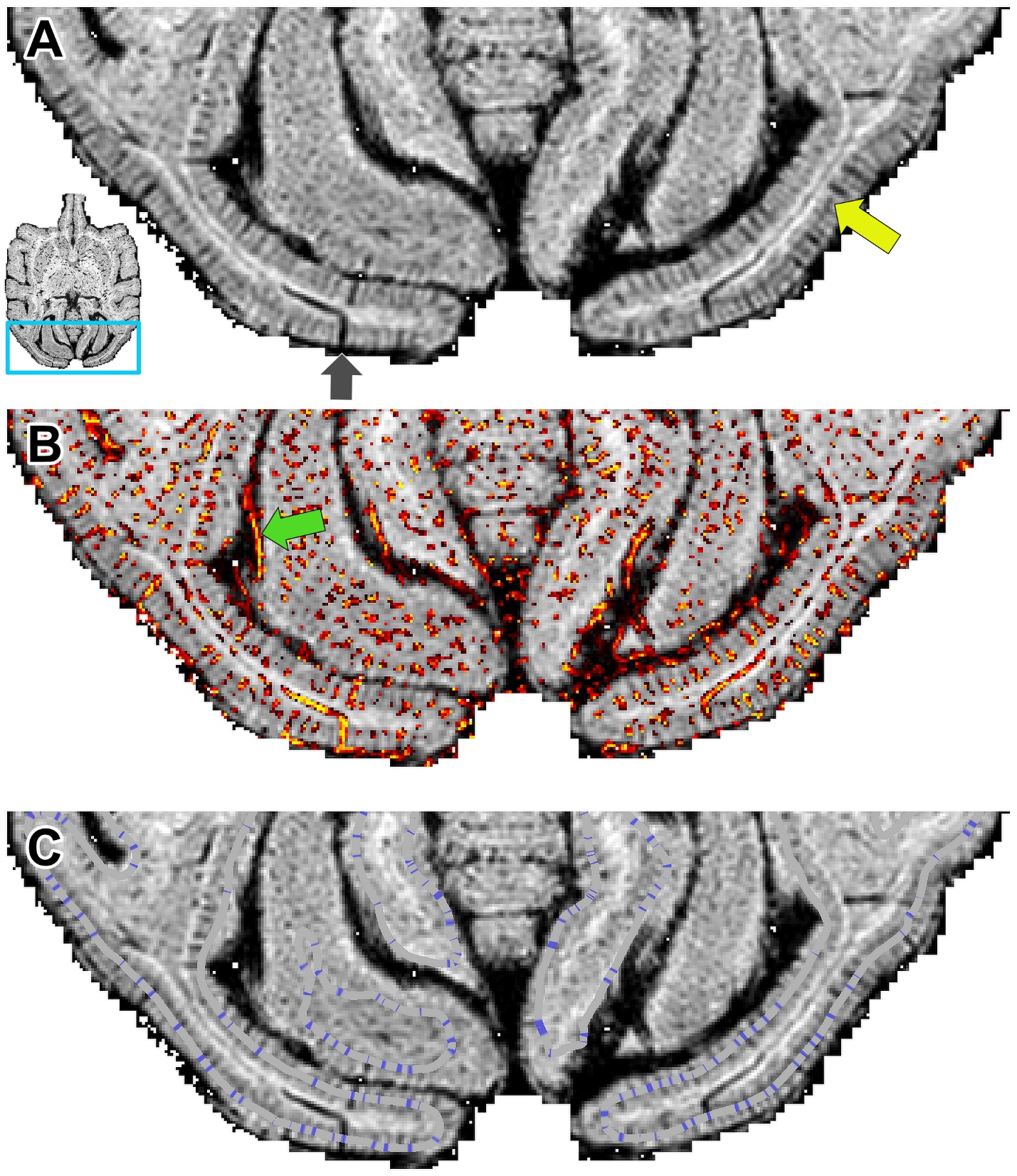

**Supplementary Figure 3. Charting vessels in the visual cortex. (A)** Ferumoxytol-weighted gradient-echo image (TE = 14 ms, 0.23 mm isotropic). The yellow arrow highlights a layer with high vascular density, likely corresponding to the primary input layer IVc. The dark gray arrow indicates a penetrating vessel extending to the white matter. Snippet shows the location of the zoomed view. **(B)** Vessel detection using Frangi filter. The green arrow denotes a pial vessel running along the cortical surface. **(C)** Vessel detection using low-frequency subtracted signal-intensity surface maps. Blue color signifies the central location of a vessel in a representative equivolumetric layer 3b. Note that not all vessels are labeled in this view as some of them are aligned orthogonal to the slice and their peaks are located in the adjacent imaging slice.

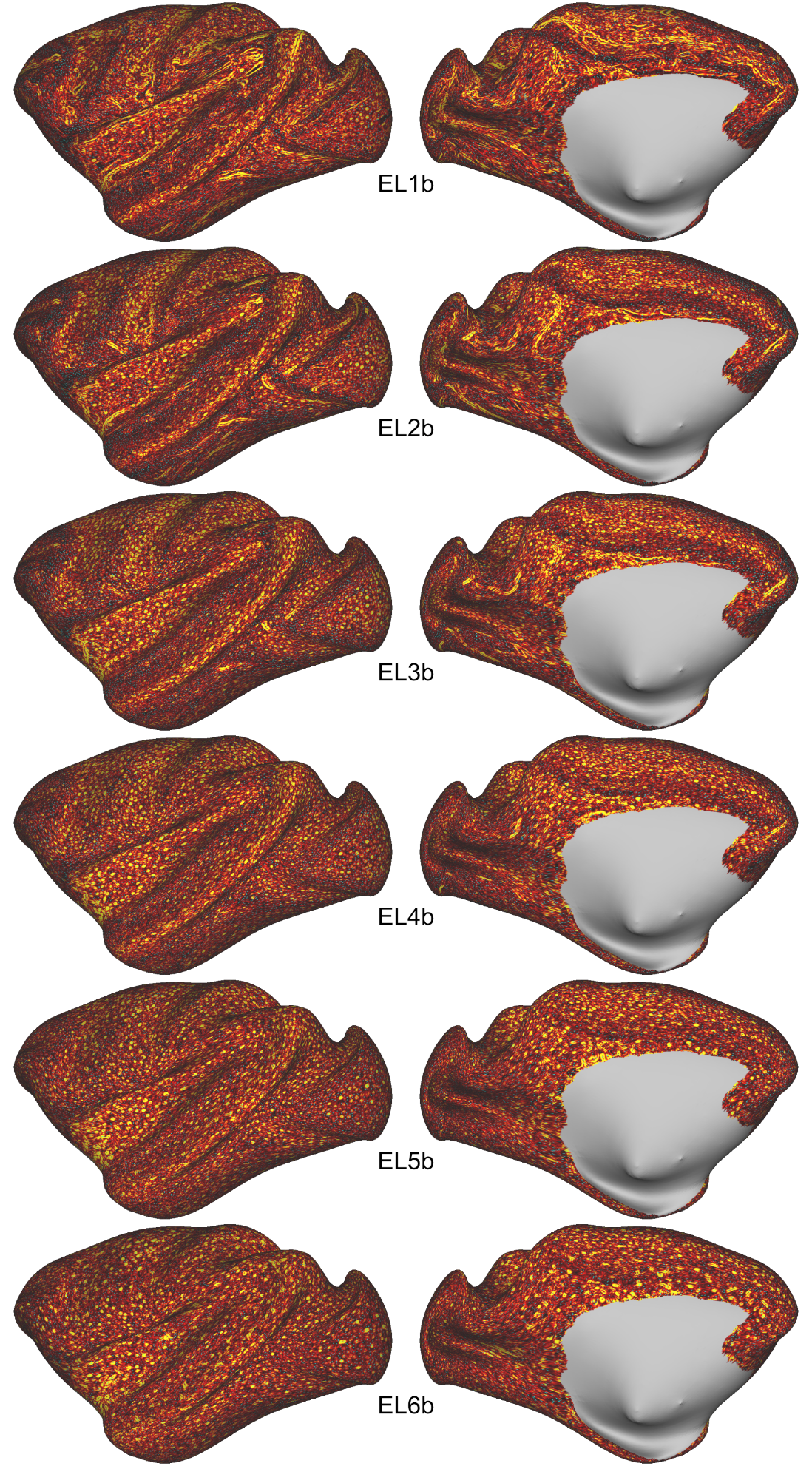

**Supplementary Figure 4.** Detection of intra-cortical vessels across equivolumetric layers.

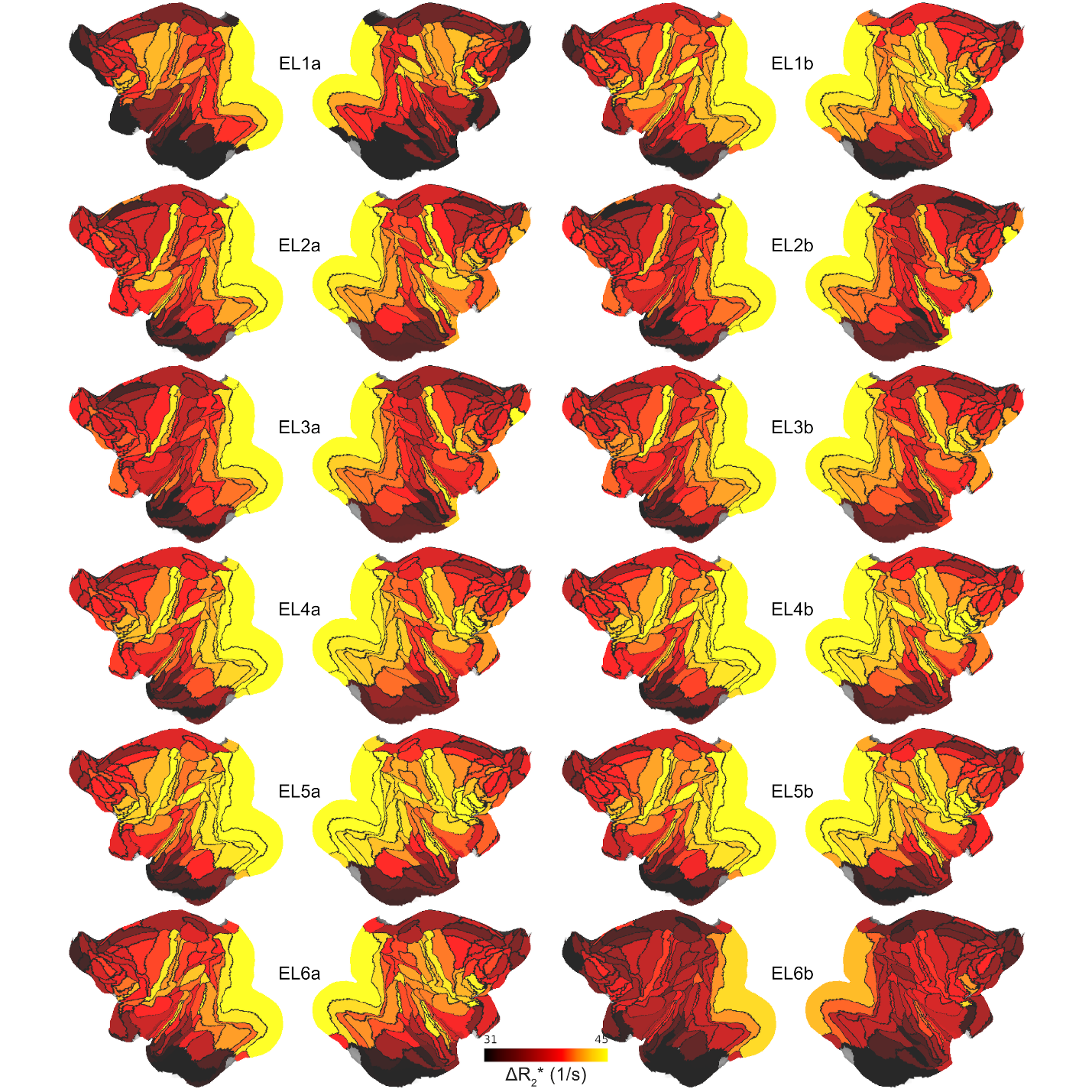

**Supplementary Figure 5. Ferumoxytol induced change in R_2_ (ΔR_2_*) across equivolumetric layers (ELs).** Data was parcellated using M132 atlas.

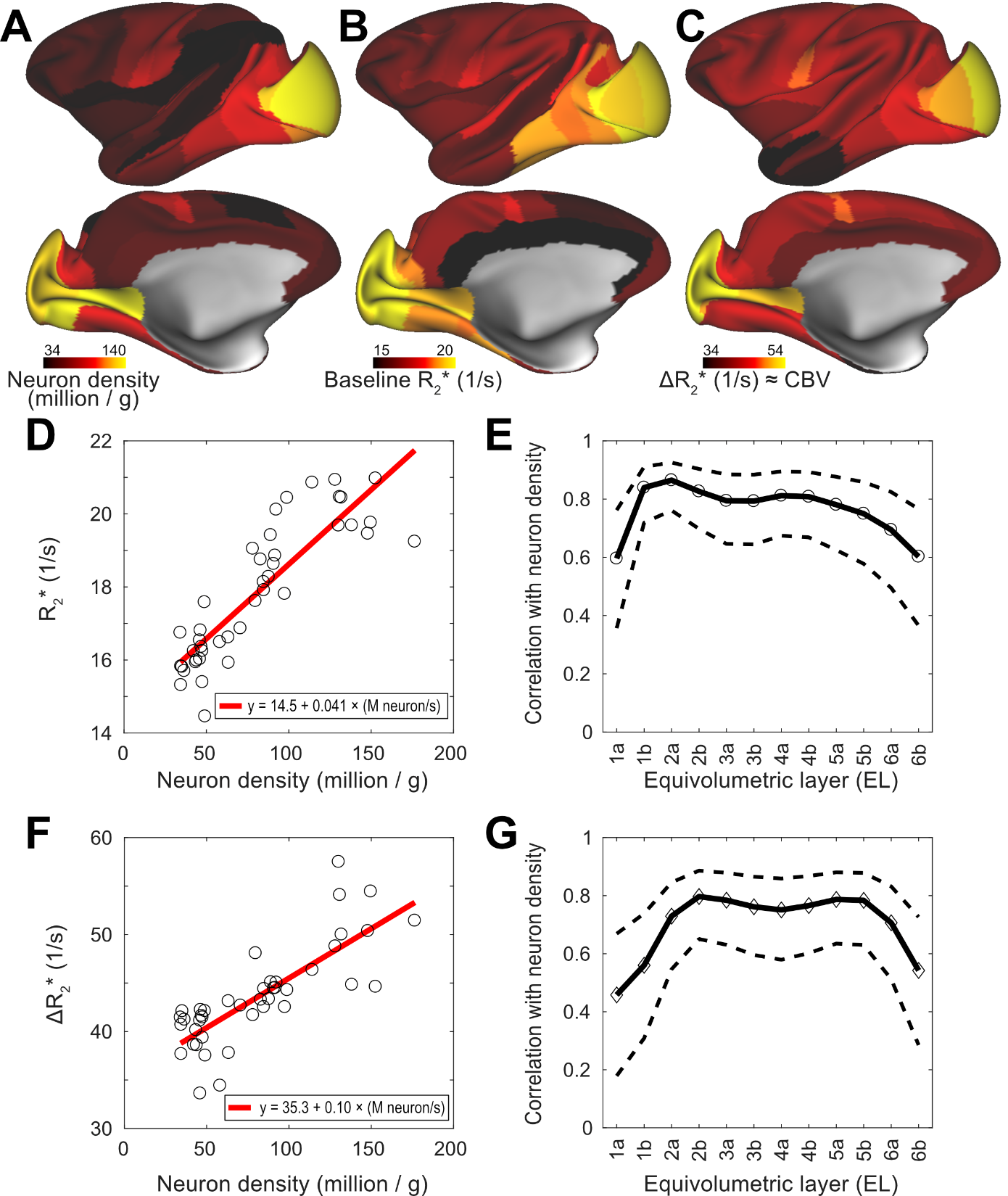

**Supplementary Figure 6. Comparison with neuron density and baseline R_2_* and ΔR_2_* in the entire cerebral cortex. (A)** Neuron density map. Data was obtained from literature (Collins et al., 2010; Froudist-Walsh et al., 2023). **(B)** Baseline R_2_* displayed in a representative equivolumetric layer 2a (EL2a) and **(C)** ferumoxytol induced ΔR_2_*, an indirect proxy measure of cerebral blood volume (CBV), displayed in EL2b. Scatter plots of **(D)** R_2_* and **(F)** ΔR_2_* plotted with respect to the neuron density in EL2a and EL2b, respectively, with red lines indicating linear fits. Pearson’s correlation coefficient between neuron density and **(E)** R_2_* and **(G)** ΔR_2_* across ELs, with dashed lines indicating 95% confidence intervals. Notably, the R_2_* intercept deviate from the free water R_2_* (≈ 1 1/s), and the ΔR_2_* intercept is non-zero, suggesting that both R_2_* and ΔR_2_* are substantially influenced by non-neuronal factors.

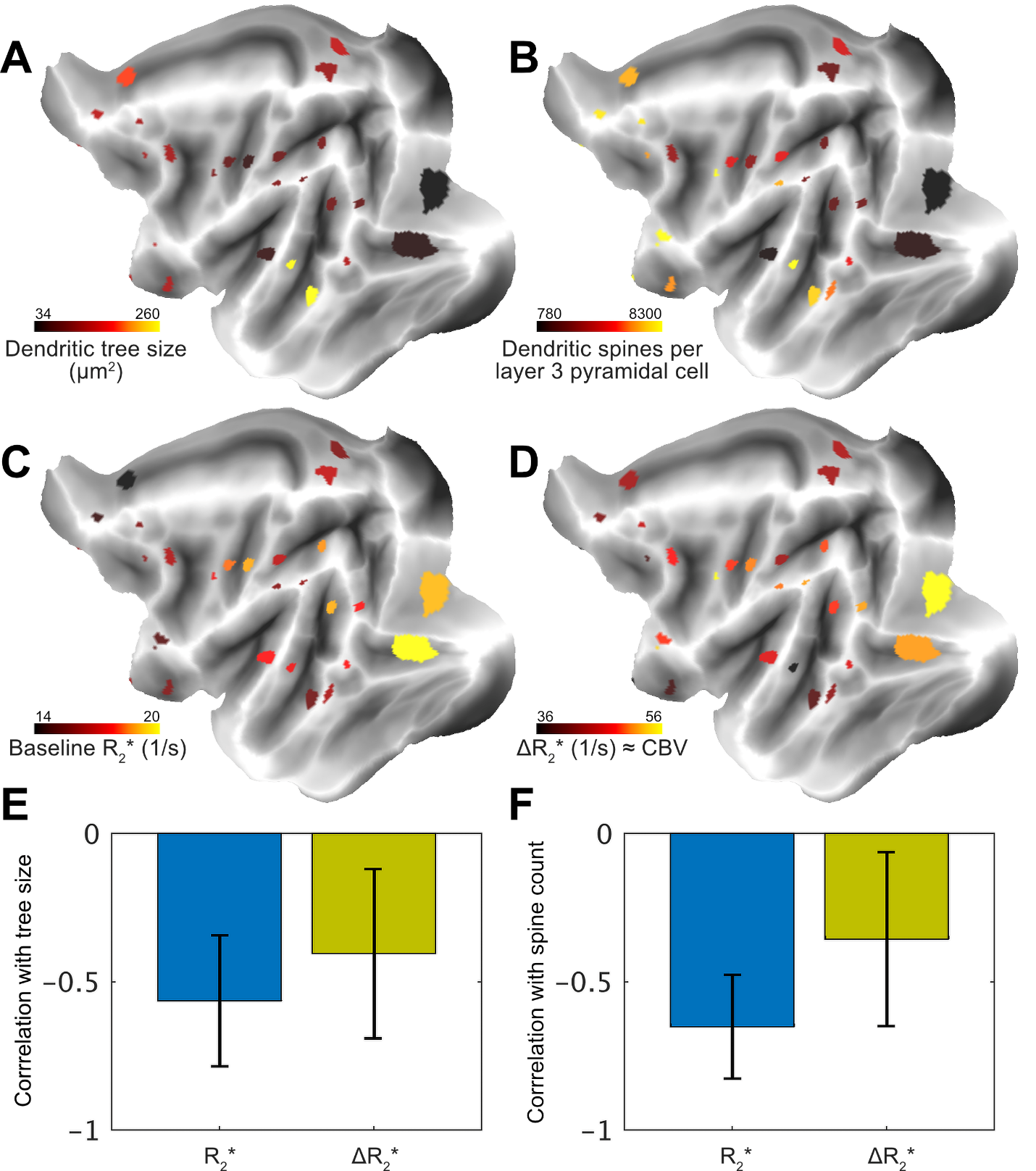

**Supplementary Figure 7. Anatomical underpinnings of heterogeneous vascular density. (A)** Dendritic tree size and **(B)** number of dendritic spines per layer 3 pyramidal cell (Elston, 2007; Froudist-Walsh et al., 2023) are compared with **(C)** baseline R_2_* and **(D)** ΔR_2_*in equivolumetric layer 3 (3a+3b). **(E)** Dendritic tree size and **(F)** spine counts are negatively correlated with cortical variation in R_2_* and ΔR_2_*. The error bars indicate 95% confidence intervals.

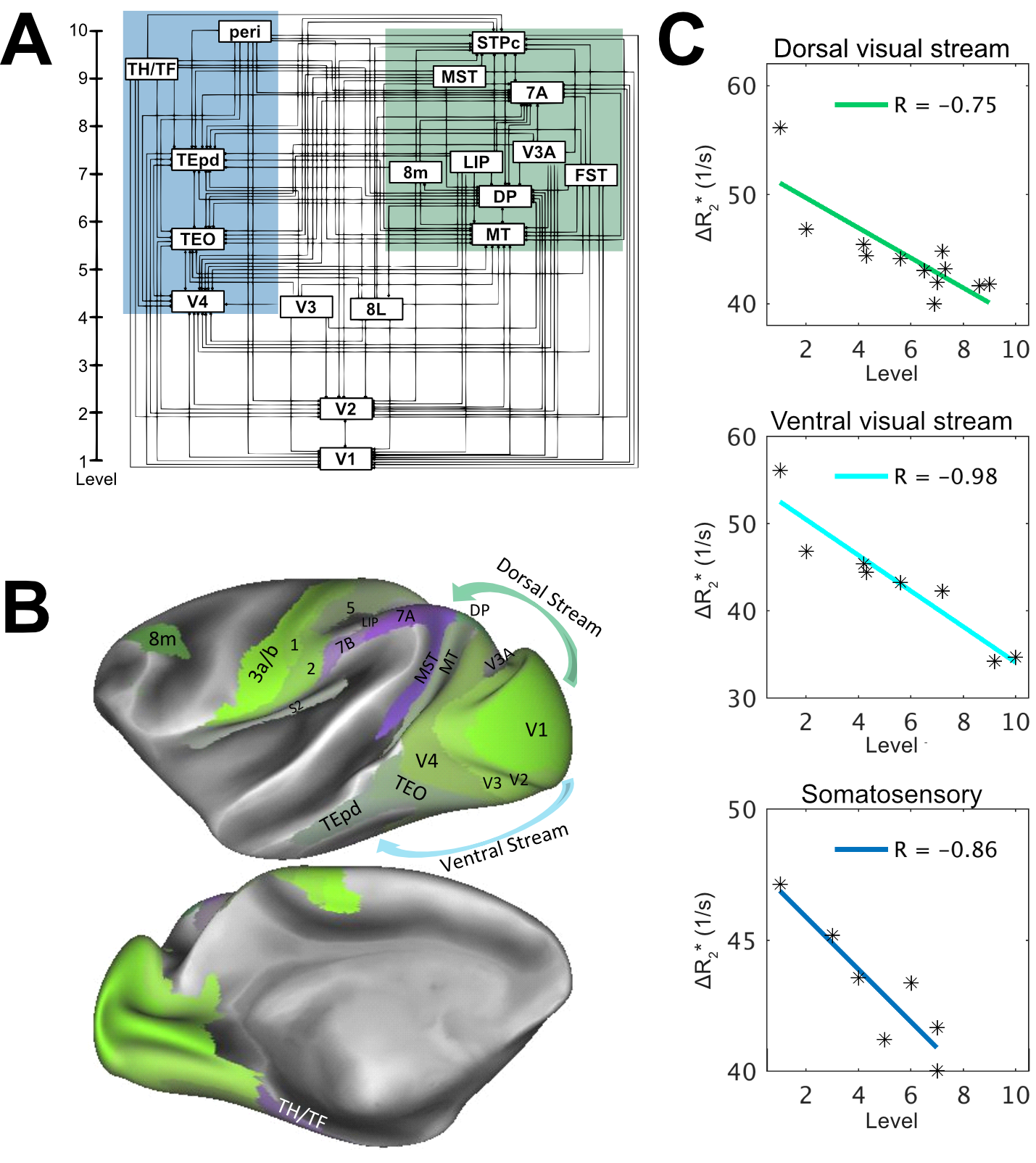

**Supplementary Figure 8. Cerebrovascular volume varies along the cortical hierarchy. (A)** Level of area within cortical hierarchy [adapted with permission (Markov et al., 2013)]. **(B)** Selected cortical areas displayed on a cortical surface map. **(C)** Ferumoxytol-induced change in transverse relaxation rate (ΔR_2_*), an indirect proxy measure of vascular volume, plotted with respect to the hierarchical level. The upper panel displays visual dorsal stream, the middle panel visual ventral stream and the bottom panel somatosensory. Hierarchical orders were obtained from (Felleman and Van Essen, 1991; Markov et al., 2014).

**References**

Collins, C.E., Airey, D.C., Young, N.A., Leitch, D.B., Kaas, J.H., 2010. Neuron densities vary across and within cortical areas in primates. Proc. Natl. Acad. Sci. 107, 15927–15932. https://doi.org/10.1073/pnas.1010356107

Elston, G.N., 2007. in Evolution of Nervous Systems (eds Kaas, J. H. & Preuss, T. M.). Elsevier, pp. 191–242.

Felleman, D.J., Van Essen, D., 1991. Distributed hierarchical processing in the primate cerebral cortex. Cereb. Cortex N. Y. N 1991 1, 1–47. https://doi.org/10.1093/cercor/1.1.1

Froudist-Walsh, S., Xu, T., Niu, M., Rapan, L., Zhao, L., Margulies, D.S., Zilles, K., Wang, X.-J., Palomero-Gallagher, N., 2023. Gradients of neurotransmitter receptor expression in the macaque cortex. Nat. Neurosci. 26, 1281–1294. https://doi.org/10.1038/s41593-023-01351-2

Markov, N.T., Ercsey-Ravasz, M., Essen, D.C.V., Knoblauch, K., Toroczkai, Z., Kennedy, H., 2013. Cortical High-Density Counterstream Architectures. Science 342. https://doi.org/10.1126/science.1238406

Markov, N.T., Vezoli, J., Chameau, P., Falchier, A., Quilodran, R., Huissoud, C., Lamy, C., Misery, P., Giroud, P., Ullman, S., Barone, P., Dehay, C., Knoblauch, K., Kennedy, H., 2014. Anatomy of hierarchy: feedforward and feedback pathways in macaque visual cortex. J. Comp. Neurol. 522, 225–259. https://doi.org/10.1002/cne.23458
